# An Integrated *In Vitro* Platform and Biophysical Modeling Approach for Studying Synaptic Transmission in Isolated Neuronal Pairs

**DOI:** 10.1101/2025.06.04.657933

**Authors:** Giulia Amos, Vaiva Vasiliauskaitė, Jens Duru, Maria Leonor Azevedo Saramago, Tim Schmid, Alexandre Suter, Ferran Cid Torren, Joël Küchler, Tobias Ruff, János Vörös, Katarina Vulić

## Abstract

Studying synaptic transmission and plasticity is facilitated in experimental systems that isolate individual neuronal connections. We developed an integrated platform combining polydimethylsiloxane (PDMS) microstructures with high-density microelectrode arrays to isolate and record single neuronal pairs from human induced pluripotent stem cell (hiPSC)-derived neurons. The system maintained hundreds of parallel neuronal pairs for over 100 days, demonstrating functional synapses through pharmacological validation and long-term potentiation studies. We coupled this platform with a biophysical Hodgkin-Huxley model and simulation-based inference to extract mechanistic parameters from electrophysiological data. The analysis of long-term potentiation stimulation using a biophysical model revealed (α-amino-3-hydroxy-5-methyl-4-isoxazole propionic acid) AMPA and (N-methyl-D-aspartate) NMDA receptor-specific alterations, providing quantitative insights into synaptic plasticity mechanisms. This integrated approach represents the first system combining isolated synaptic pairs, long-term stability, and mechanistic modeling, offering unprecedented opportunities for studying human synaptic function and plasticity.

## Introduction

The brain”s ability to learn, adapt, and perform complex functions arises from the dynamic interplay of neuronal networks that selectively transmit and store information. Learning occurs when experience induces lasting changes in neuronal activity and behavior. At the cellular level, these adaptive changes are mediated primarily by synaptic plasticity, defined as the ability of synapses to strengthen or weaken over time in response to patterns of activity. Understanding how plasticity shapes the connectivity and function of neuronal networks is central to understanding the mechanisms underlying learning, memory, and brain development. Research in this area follows two complementary avenues: experimental approaches and computational modeling.

Traditional experimental approaches to studying synaptic function and plasticity span a spectrum from complex *in vivo* and *ex vivo* preparations to simplified and highly controlled *in vitro* systems. *In vivo* approaches have revealed fundamental principles such as long-term potentiation (LTP) and long-term depression (LTD), and have mapped the connectivity and plasticity rules of various brain regions [1, 2, 3, 4]. Brain slices, for example, preserve local circuit architecture and have enabled detailed characterization of synaptic responses and plastic changes following stimulation protocols [5, 6, 7, 8]. However, these experimental models present two significant limitations: first, they rely on animal preparations rather than human tissue, potentially limiting translational relevance; and second, they cannot effectively isolate the contributions of individual synapses or neuronal pairs due to extensive background connectivity and overlapping activity from surrounding neuronal networks. Consequently, unraveling the precise cellular and molecular mechanisms underlying synaptic transmission and plasticity remains challenging in these complex preparations.

*In vitro* experiments offer compelling solutions to these challenges by providing greater experimental control and human relevance. First, they enable the use of human-derived cells through induced pluripotent stem cell (iPSC)-derived neurons, eliminating species-specific differences that may limit translational applicability. Second, they provide simplified systems with substantially fewer confounding variables than intact brain preparations [9, 10, 11, 12]. However, even random *in vitro* cultures, despite containing orders of magnitude fewer neurons than the brain, can develop complex network architectures that obscure fundamental mechanisms. Therefore, imposing structural organization and directional connectivity in cultured systems becomes crucial for mechanistic studies. One promising approach involves guiding neuronal growth and controlling connectivity by confining cells within polydimethylsiloxane (PDMS) microstructures. Substantial progress has been made in this direction, ranging from larger compartmentalized systems that isolate neuronal clusters [13, 14, 15] to progressively smaller configurations designed to isolate individual neurons[16] and axons [17, 18, 19] or even synaptic locations [20, 21]. While experimental approaches provide empirical data on synaptic function [11, 12], computational models complement these efforts by allowing researchers to manipulate parameters and test hypotheses that would be challenging to address experimentally [22, 23]. Computational models have elucidated the rules governing spike-timing-dependent plasticity [24, 25, 26], modeled the impact of synaptic noise and variability [27], and predicted how changes at individual synapses can scale up to affect network dynamics and information processing[28, 29]. These types of models play an important role in bridging the gap between molecular mechanisms and emergent network behavior. However, despite their power, computational models are inherently limited by the need to simplify biological detail for tractability [23, 30, 31], and their predictions often lack direct experimental validation due to difficulties in finding precise biological equivalents of the computational networks. This gap in the experimental validation highlights the need for experimental systems that can serve as a bridge between computational models and biological approaches. Small and well-characterized *in vitro* experimental systems represent ideal candidates for biophysical modeling approaches [32, 33]. Networks with reduced complexity allow for a comprehensive parametrization of synaptic properties without the confounding influences present in more complex systems. The reduction in variables, improved experimental control, and clearer correspondence between model components and biological systems enable more direct comparison of experimental observations with model predictions, facilitating rigorous validation of computational frameworks against empirical data. Consequently, well-characterized small systems provide optimal platforms for iterative model refinement, where discrepancies between predicted and observed behavior can be systematically addressed through gradually extending the model with missing details. Model predictions can then guide experimental investigations, establishing a bidirectional feedback loop that accelerates both theoretical understanding and empirical hypothesis testing. Fully realizing this integrative approach requires appropriate measurement technologies that balance resolution with throughput. Patch clamp electrophysiology has been the gold standard for obtaining precise information about ionic channel properties, synaptic currents, and membrane dynamics, providing the detailed biophysical data necessary for model parametrization and validation. However, while patch clamping offers unparalleled resolution for studying individual synapses and neurons, these approaches are inherently time-consuming, low-throughput, and incompatible with long-term monitoring. Therefore, higher-throughput recording methods, such as extracellular microelectrode arrays (MEAs) are essential to generate statistically robust datasets while, to some extent, maintaining the cellular-level resolution required for detailed modeling [34, 9].

In this study, we present a platform that integrates experimental and computational models with compartmentalized PDMS microstructures for studying synaptic function. We aim to isolate the simplest network-level functional unit of neuronal communication, single neuronal pairs, to create a basic system for studying unidirectional synaptic transmission. By employing PDMS microstructures with precisely defined dimensional constraints, we enable stochastic isolation of ultra-low-density networks, down to individual neuronal pairs. The microscale dimensions of these channels allow hundreds of isolated pairs to be positioned adjacently on a single device. Furthermore, the compatibility of the microstructures with various substrates, particularly high-density microelectrode arrays, enables parallel readout capabilities that provide statistically robust large datasets while maintaining access to individual neuronal dynamics over several months. This combination of isolation, parallelization, and long-term stability creates opportunities for detailed investigation of synaptic mechanisms based on extracellular electrophysiology in hiPSC-derived neuronal networks.

We demonstrate the capability of this integrated platform to characterize isolated functional neuronal pairs derived from human iPSC-derived excitatory neurons. We showcase the versatility of the PDMS microstructure designs, presenting multiple geometric configurations that can be tailored for diverse experimental applications, from basic connectivity studies to modulating the neuronal dynamics. By developing a comprehensive biophysical model based on Hodgkin-Huxley formalism [35, 33], we show the feasibility of combining our simplified experimental system and biophysical computational modeling to effectively bridge empirical observations with mechanistic insights. Leveraging simulation-based inference techniques [36, 37], we use this model to quantitatively characterize synaptic transmission properties and extract biophysically meaningful parameters from the experimental data. As a proof-of-concept application, we implemented a well-established LTP stimulation protocol and analyzed the resulting changes in model parameter distributions, revealing AMPA and NMDA receptor-specific alterations that provide mechanistic insights into synaptic plasticity. Together, these results establish this platform as a powerful tool for investigating fundamental questions about human synaptic function and plasticity with unprecedented precision and throughput.

## Star Methods

Human induced pluripotent stem cell (hiPSC) derived neurogenin-2 (NGN2) excitatory neurons (hereafter referred to as iNeurons) were cultured inside single-cell polydimethylsiloxane (PDMS) microstructures either on top of glass substrates for visual assessment and quantification or on top of HD-CMOS MEAs for electrophysiology characterization.

### PDMS microstructures

PDMS microstructures for isolating and guiding single cells were designed using CAD software (AutoCAD 2021) and fabricated by Wunderlichips (Zurich, Switzerland). More details on the fabrication process are available in previous publications [?, 21, 18]. The PDMS microstructures were fabricated with openings of 10 µm (see schematic in Fig. 1A). The microchannels were designed with two distinct heights: 2 µm for structures containing submicrometer features and 4 µm for those without, while maintaining a consistent width of 10 µm across all channels (see schematic in Fig. 1A,B). The total thickness of the PDMS microstructures was 75 µm. To accommodate axonal growth, the microchannels were designed with variable lengths, all exceeding 2.5 mm, and terminated in a loop structure to entrap the extending axons. Each well was optimized to house a single neuron, with the total number of wells varied based on the specific application requirements. A photo of the PDMS membrane with visible channel and well contours is shown in Fig. 1Aiii.

**Fig. 1:**
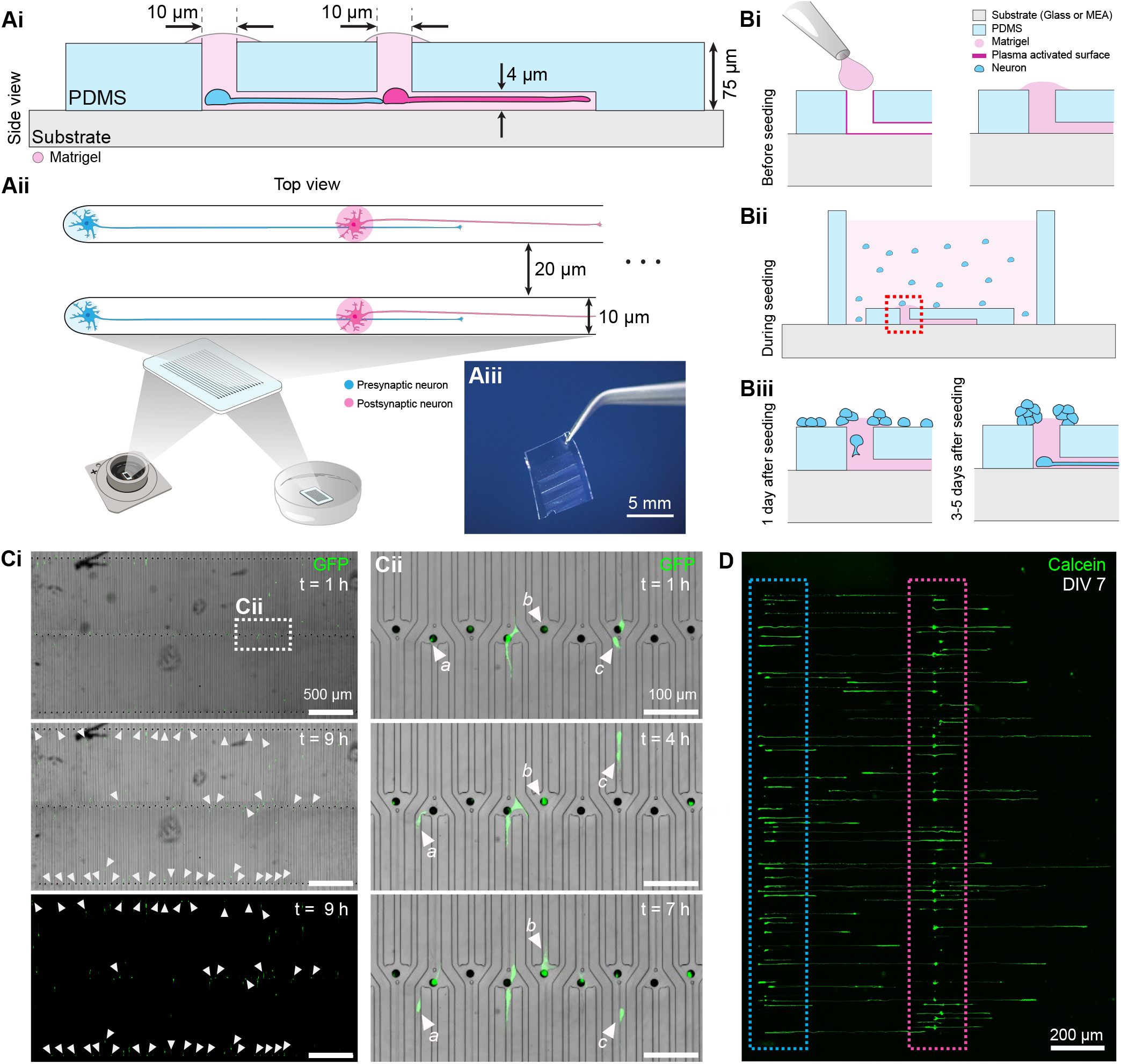
Single-cell PDMS microstructure for neuronal isolation and migration. A) Device design concept and adaptability to substrates. Ai) The PDMS microstructure features 10 µm wide and 75 µm high wells that narrow down to 4 µm high microchannels. Microstructures are placed on a substrate and filled with matrigel. Aii) The microchannel design connects two wells intended for presynaptic (blue) and postsynaptic (pink) neurons. Microchannels isolating separate pairs are min. 20 µm apart. Aiii) Photograph showing an actual fabricated PDMS device as used in our experiments. B) The cell seeding and migration process. Bi) Prior to cell seeding, the microchannels are filled with matrigel to create a supportive matrix for cell growth. Bii) During seeding, neurons randomly position themselves on top of the microstructure. 0.5 mm high PDMS seeding frame is placed around the microstucture to restrict the area where the cells can land. Biii) Following seeding, neurons gradually migrate into the microchannels, with movement observed after one day and continued migration within the first week. C) Timelapse of neuronal migration upon seeding. Ci) Phase contrast images reveal the progressive migration of cells into the microwells during the first hour (t = 1 h) and after 9 hours (t = 9 h) with GFP-labeled neurons. Neurons that enter the microchannels between the first and ninth hour are indicated with white arrows. Cii) Higher magnification views of the boxed region allow for tracking individual cells (three exemplary cells marked by letters) as they navigate through the microchannels. D) Calcein-stained neurons on DIV 7 grew along a presynaptic (indicated with a blue rectangle) and postsynaptic (indicated with a pink rectangle) region.

### Substrate preparation and surface functionalization

The PDMS microstructures were placed either on top of glass substrates (35 µm Ibidi, 81218-200-IBI, Vitaris, Switzerland) for visual assessment and quantification or on top of high-density complementary metal oxide semiconductor microelectrode arrays (HD-CMOS MEAs) for electrophysiology characterization. Before mounting, both the microstructure and the substrate were cleaned and activated in a Tergeo Plasma Cleaner (Pie Scientific, USA) using an O_2_ + H_2_O mixture for 1 minute. For CMOS chips, care was taken to leave a portion of the reference electrode uncovered during mounting. After a 2-minute waiting period to ensure initial attachment, the surfaces were functionalized with matrigel (Corning matrigel Basement Membrane Matrix, 354234, 10.5 mg/ml) diluted to 3 mg/ml in neurobasal medium. The matrigel solution was prepared 1-12 hours before use and maintained at 4°C to prevent premature crosslinking. Approximately 2.5 µL of the cold matrigel solution was carefully dispensed over the microstructure openings and guided through the microchannels via gentle pipetting. Excess matrigel was removed by aspiration, and the substrates were incubated at 37°C for 5 minutes. To minimize overgrowth, an optional second plasma treatment (2 minutes, O_2_ + H_2_O) was performed to selectively remove matrigel from the top surface of the microstructure. The devices were then filled with phosphate-buffered saline (PBS) and desiccated for approximately 5 minutes to remove any air bubbles. Finally, the PBS was replaced with 500 µL of complete neurobasal medium.

### Cell culture

The following types of hiPSC-derived NGN2 neurons were used: green fluorescent protein (GFP) expressing NGN2 neurons and unlabeled NGN2 neurons.

### hiPSC differentiation

NGN2 neurons were generated following Giorgetti et al. protocol [38] and transfected with doxycycline-inducible NGN2. Neuronal differentiation was induced by 3-day doxycycline exposure. Differentiated neurons were cryopreserved in aliquots of 1-8*·*10^6^ cells in FBS with 5% DMSO, provided by Novartis and stored in liquid nitrogen until use. More details are available in our previous work [39].

### hiPSC neuron seeding and maintenance

An aliquot containing iNeurons was removed from liquid nitrogen storage and rapidly thawed at 37°C. The 1 mL thawed cell solution was transferred dropwise into 4 mL of warm NBD and centrifuged for 5 minutes at 1000 rpm. After aspirating the supernatant, cells were resuspended to a concentration of 1*·*10^6^ cells per mL. The substrate was retrieved from the incubator PBS was aspirated and a sterilized PDMS frame (1×1×1 cm^3^) was positioned on top of the microstructure. Approximately 50 µL of warm medium was pipetted into the frame. The volume containing 50,000 cells was then pipetted into the frame and the dish was placed in the incubator. After approximately 20 minutes, an additional 500 µL of warm medium was added around the frame to prevent excessive evaporation. The frame was removed the following day, and the medium was exchanged and topped up to approximately 1 mL.

The cells were cultured in NeuroBasal medium (NB) (21203-049, ThermoFisher) [40]. Fresh NB complete medium was prepared, consisting of a 2% solution of B-27 supplement (17504-044), a 1% solution of Penicillin-Streptomycin (P-S) (15070-063), and a 1% solution of GlutaMAX (35050-061), all sourced from ThermoFisher. In the first 10 days, doxycycline (Clontech Labs 3P 631311, Fisher Scientific) was added in a 1:500 ratio to further enhance neuronal differentiation and maturation.

### Staining and imaging

iNeurons were imaged on day in vitro (DIV) 7 as live cultures to assess channel occupancy and culture viability. Following neuronal maturation and synapse formation, selected samples were subsequently fixed, immunolabeled, and visualized for further analysis.

### Live and dead cell staining

The growth of cells in single-cell PDMS microstructures was monitored using NeuroFluor™ NeuO (01801, StemCell Technologies, Switzerland). NeuO is a membrane-permeable fluorescent probe that selectively labels primary and iPSC-derived neurons. On the day of imaging, cells were stained following the standard protocol provided by the manufacturer. Some cultures were stained with Calcein and Ethidium Homodimer at a 1:1000 ratio.

### Immunofluorescence staining

The samples were fixed in the fourth or fifth week in culture using 4% paraformaldehyde (1004960700, Sigma-Aldrich) for 15 min at room temperature. Following fixation, samples were washed three times with PBS, allowing 10 min intervals between washes. Cell membranes were then permeabilized using PBS containing 0.1% Triton X-100 (X100, Sigma-Aldrich) for 5-8 minutes, after which the permeabilization solution was removed by three consecutive PBS washes with appropriate waiting periods. Subsequently, samples were incubated with primary antibody solution containing goat serum (31873, Invitrogen) and the following primary antibodies at 1:1000 dilution: rabbit anti-Synapsin (AB254349, Abcam), mouse anti-PSD-95 (AB13552, Abcam), and chicken anti-neurofilament (AB4680, Abcam). To enhance antibody diffusion into the PDMS microchannels, samples were incubated for 48 h on a shaker at room temperature. Following primary antibody incubation, samples were washed three times with PBS (10 min per wash) and subsequently incubated with secondary antibody solution consisting of Alexa Fluor 488 goat anti-chicken (A11039, Life Technologies), Alexa Fluor 555 goat anti-rabbit (A32732, Invitrogen), and Alexa Fluor 647 goat anti-mouse (A32728, Invitrogen) antibodies diluted 1:500 in goat serum. Secondary antibody incubation was also performed for 48 h on a shaker at room temperature. After incubation, samples were washed three times with PBS (10 min intervals) and counterstained with Hoechst (1:500 in PBS) for approximately 1 h. Finally, samples underwent three additional PBS washes and were stored at 4°C until imaging, which was performed either on the same or the following day.

### Image acquisition and analysis

Cells cultured in PDMS microstructures on glass substrates were visualized using a confocal laser scanning microscope (CLSM) (Fluoview 3000, Olympus). For live cell imaging, either a 10× (Olympus, UPLFLN10×2PH, NA = 0.3) or 20× (Olympus, UPLFLN20XPH, NA = 0.5) objective was employed. To enable visualization of synaptic puncta in immunostained samples, imaging was performed using a 60× oil-immersion objective (Olympus, UPLSAP060XS2, NA = 1.3).

Microscope images were analyzed using Fiji software [41]. To improve axonal visibility relative to soma, a pixel logarithm transformation was applied to all representative fluorescent images presented in this paper, with the exception of immunofluorescent staining images. Background fluorescence was minimized through manual adjustment of brightness and contrast parameters.

## Electrophysiology

### Microelectrode arrays for recording and modulating neuronal activity

The experiments utilized high-density complementary metal-oxide-semiconductor (HD-CMOS) microelectrode arrays (MEAs) from Maxwell Biosystems (Switzerland). These chips (MaxOne+ Chip (uncoated Pt-electrodes)) feature a flat surface topology and incorporate 26,400 electrodes arranged across a 3.85 × 2.10 mm^2^ area, with electrodes spaced at 17.5 µm intervals. Through a switch matrix, any 1,020 electrodes can be selected for simultaneous recording, with data acquisition performed at 20 kHz. The system also enables extracellular stimulation through the same electrodes using 32 independent stimulation buffers.

### Visualization of PDMS microstructure with a CMOS MEA

A custom Python script was used to generate a voltage map displaying the positions of electrodes beneath the PDMS microstructure. An example of this voltage map for the PDMS microstructure is illustrated in Fig. S13. This mapping enables precise identification and selection of electrodes that correspond to individual networks within the PDMS microstructure. A detailed description of this methodology can be found in our previous publication [42].

### Recording area selection

Upon first recording, the total chip area was split into separate electrode subselections that contained no more than 1,020 electrodes (upper limit for the number of electrodes recorded simultaneously). This number ranged from 6 to 11 areas, depending on the voltage map (PDMS placement quality).

### Data collection

The combination of subselections of the recording area with PDMS microstructures facilitated repeated recordings of identical areas, and consequently the same neurons, over time. For recording sessions, MEA chips were inserted into the recording units, which were placed in an incubator maintained at 35 °C, 5 % CO_2_, and 90 % humidity. Following placement of the culture in the incubator, a 5-minute settling period was implemented to allow CO_2_ levels and humidity to stabilize after door opening and to permit culture equilibration. Spontaneous neuronal activity was subsequently recorded for 2-5 minutes per subselected area at a sampling frequency of 20 kHz. Recordings were conducted approximately weekly during the first five weeks in culture and less frequently at later time points, with the most extended recording occurring at DIV 234 (Figure S14).

### Chemical Suppression of Synapses

Cultures at DIV 100 were treated with the synaptic blockers (6-nitro-7-sulfamoylbenzo(f)quinoxaline-2,3-dione) NBQX and AP5, which act as selective antagonists for AMPA and NMDA receptors, respectively. Each antagonist was diluted in NBD at a ration 1:1000. Spontaneous activity was first recorded during two consecutive 2-minute periods for each selected chip area to establish baseline measurements. Following baseline recording, both blockers were added to the culture medium, and spontaneous activity was immediately measured for three consecutive 2-minute periods in each area to capture the acute effects of synaptic inhibition. The culture medium was then aspirated and replaced with fresh NBD medium three times to ensure complete washout of the synaptic blockers. To assess potential long-term effects, spontaneous activity was recorded again for 2 minutes in each area one week after treatment.

### Stimulation Paradigm

Stimulation experiments shown in Figure 8 were performed on DIV 32, 35, 41, 46. The stimulation paradigm consisted of 5-min spontaneous recording which was followed by a 5-min stimulation. Stimulation consisted of 5-second 1.5 V peak-to-peak stimulation pulses (biphasic, first pulse negative, total pulse duration 400 µs) delivered at a 5 Hz frequency, interchanged with 5 second break time. Two 5-min spontaneous recordings were performed sequentially after stimulation.

### Data analysis: neuronal pair identification and synaptic pair calculation

Data was processed and analyzed using a custom-made data analysis pipeline. First, spikes were detected and sorted using the Spikeinterface framework [43] including SpyKING CIRCUS v2 spike sorter [44]. The spike detection parameters included detection at a negative peak exceeding 5 times the signal standard deviation and 3 ms minimum spike distance. Identified spike units were excluded if the signal-to-noise ratio was below 5 or the firing rate of the spike unit was below 0.1 Hz. The following metrics were calculated per unit using custom-made scripts: firing rate, conduction speed, and waveform metrics: amplitude, peak-trough ratio and duration, half width, repolarization slope and recovery slope.

To identify putative synaptic pairs *i.e*., synaptic coupling between units, multivariate transfer entropy (mTE) was used. mTE is an information-theoretic measure that quantifies the directed transfer of information between multivariate time series. It extends the concept of transfer entropy by accounting for the influence of multiple variables on the information flow. Consider two stochastic processes X and Y. Each stochastic process is a collection of random variables {*X*_*t*_}, where for each variable its realized value is denoted by *x*_*t*_. We will also use length-*𝓁* and length-*k* collections of random variables (embedding vectors) 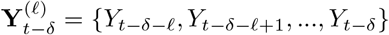 and 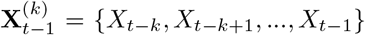, where *δ* is the time delay. Transfer entropy from Y to X is defined as the reduction in uncertainty about the future state of X when considering the past states of Y, over and above what is already available in the past states of Xbossomaier2016transfer,paluvs2001synchronization, schreiber2000measuring expressed in the form of conditional mutual information:

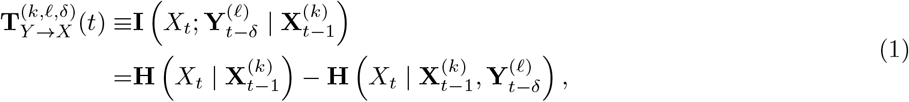

where *I*(*X*; *Y*) is mutual information between the two random variables *X* and *Y* that quantifies the reduction in uncertainty (entropy) about one variable given knowledge of the other.

mTE generalizes Eq. 1 to account for the influence of other potential sources. This is crucial for distinguishing direct from indirect information transfer, especially in networks where multiple units may be interconnected or driven by common inputs. mTE from Y to X is conditioned on a collection of other stochastic processes Z:

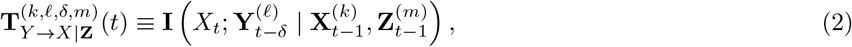

where 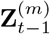 denotes the past of the conditioning set (collection of stochastic processes) and *m* is the corresponding embedding length.

Inference of the functional connectivity using mTE was done using IDTxl [45]. The IDTxl software utilizes an optimization approach that combines greedy search algorithms with comprehensive null hypothesis testing to determine the optimal set of parent nodes Z and historical embeddings for both the past of X and Y.

Despite the optimization, computational demands for mTE calculation remain large: complete network analysis has a computational complexity of *O*(*N*^2^ *× d × τ*_max_ *× S*), where *N* represents neuron count, *d* indicates average incoming connections per neuron, *τ*_max_ denotes maximum temporal search depth, and *S* represents the number of surrogate tests [46, 47]. Due to these computational constraints, a preselection step was implemented to reduce the number of potential incoming connections to each neuron as previously described in Varley et al [46]. The incoming source nodes were prefiltered for each target neuron by only including neurons that demonstrated statistically significant bivariate transfer entropy (Eq. 1) to the target neuron across temporal search depth *τ*_max_ of 10 ms for the source and 5 ms for the target. Statistical significance was evaluated using the empirical null hypothesis test, implemented in JIDT software package [48]. The null distribution was inferred using 200 permutations. False discovery rates were corrected for using the Benjamini-Hochberg correction for multiple testing [49]. Significant source-target pairs as identified with bivariate transfer entropy were used to compute multivariate transfer entropy (Eq. 2) using IDTxl. Following other work on neuronal data [46, 50, 51], only one bin of source history was considered, defined as the lag that maximized the significant bivariate transfer entropy. On the other hand, the target history was searched using a search depth *τ*_max_ of 10 ms.

Once synaptic pairs were identified, defined as source and target neurons with a significant multivariate transfer entropy (p *≤* 0.05), the probability of postsynaptic spike was calculated as follows:

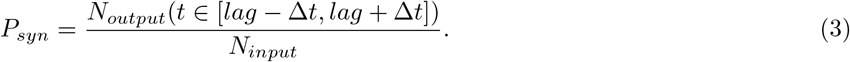

Pairs with a postsynaptic spike probability greater than 0.05 were kept for further analysis.

### Computational Model

*In silico*, a network is modeled as two synaptically connected multicompartmental neurons, as well as extracellular electrodes that capture the electrical field generated by the traveling action potentials. The model is implemented using LFPy library [52], the model of a neuron and a synapse are defined using NEURON simulator [53]. The model”s parameters and variables are summarized in Table 1.

**Table 1:** Fixed and variable model parameters.

| Parameter | Value / Range | Type | Explanation |
| --- | --- | --- | --- |
| **Biophysical Properties** |  |  |  |
| $r_a$ | $100\ \Omega \cdot \text{cm}$ | Fixed | Specific axial resistance |
| $c_m$ | $1\ \mu\text{F}/\text{cm}^2$ | Fixed | Specific membrane capacitance |
| $g_{Na}$ | $0.05\ \text{S}/\text{cm}^2$ | Fixed | Maximal sodium conductance |
| $g_K$ | $0.005\ \text{S}/\text{cm}^2$ | Fixed | Maximal potassium conductance |
| $g_L$ | $0.0003\ \text{S}/\text{cm}^2$ | Fixed | Leak conductance |
| $T$ | $310\ \text{K}\ (37^\circ\text{C})$ | Fixed | Temperature |
| **Reversal Potentials** |  |  |  |
| $E_L$ | $-39.2\ \text{mV}$ | Fixed | Leak reversal potential |
| $E_{Na}$ | $70\ \text{mV}$ | Fixed | Sodium reversal potential |
| $E_K$ | $-80\ \text{mV}$ | Fixed | Potassium reversal potential |
| $V_T$ | $-30.4\ \text{mV}$ | Fixed | Threshold shift for sodium activation |
| **Morphology** |  |  |  |
| $d_{\text{soma}}$ | $10\ \mu\text{m}$ | Fixed | Soma diameter |
| $\ell_{\text{soma}}$ | $10\ \mu\text{m}$ | Fixed | Soma length |
| $\ell_{\text{axon}}$ | $2000\ \mu\text{m}$ | Fixed | Axon length |
| $d_{\text{axon}}^{\text{pre}}, d_{\text{axon}}^{\text{post}}$ | $[0.2, 5]\ \mu\text{m}$ | Variable | Axon diameters of pre- and post-synaptic neurons |
| **Electrode Geometry** |  |  |  |
| $(x_{e1}, y_{e1}, z_{e1})$ | $(0, 0, 60)\ \mu\text{m}$ | Fixed | Electrode 1 position |
| $(x_{e2}, y_{e2}, z_{e2})$ | $(0, 0, 1060)\ \mu\text{m}$ | Fixed | Electrode 2 position |
| **Synaptic Properties** |  |  |  |
| $z_{\text{syn}}$ | $[1000, 2000]\ \mu\text{m}$ | Variable | Synapse coordinate |
| $\mu_{\text{AMPA}}$ | $[0.0, 0.002]$ | Variable | Mean value of AMPA synaptic weight |
| $\sigma_{\text{AMPA}}$ | $[0.0, 0.001]$ | Variable | Standard deviation of AMPA synaptic weight |
| $\mu_{\text{NMDA}}$ | $[0.0, 0.0002]$ | Variable | Mean value of NMDA synaptic weight |
| $\sigma_{\text{NMDA}}$ | $[0.0, 0.0001]$ | Variable | Standard deviation of NMDA synaptic weight |
| $\tau_2^{\text{AMPA}}, \tau_1^{\text{AMPA}}, e_{\text{AMPA}}$ | $(0.1\ \text{ms}, 0.3\ \text{ms}, 0\ \text{mV})$ | Fixed | Timescales and reversal potential of AMPA receptors |
| $\tau_2^{\text{NMDA}}, \tau_1^{\text{NMDA}}, e_{\text{NMDA}}$ | $(10\ \text{ms}, 30\ \text{ms}, 0\ \text{mV})$ | Fixed | Timescales and reversal potential of NMDA receptors |
| **Spontaneous Input Current $I_{\text{spont}}$** |  |  |  |
| $\lambda_{\text{pre}}, \lambda_{\text{post}}$ | $[5, 100]\ \text{Hz}$ | Variable | Poisson process rate |
| $\tau_1, \tau_2, e$ | $0.2\text{ms}, 0.4\text{ms}, 0\text{mV}$ | Fixed | Timescales and reversal potential of $I_{\text{spont}}$ |
| $\mu_{\text{spont}}$ | $0.02$ | Fixed | Mean value of spontaneous current |
| $\sigma_{\text{spont}}$ | $0.02$ | Fixed | Standard deviation of spontaneous current |

#### Morphology

As stated before, a single neuron is modeled using multicompartmental formalism. The morphology is of a simple ball-and-stick neuron, with two compartments: a soma and an axon. The soma is comprised of a single segment, modeled as a cylinder with the height *h* and the diameter *d* of 10µ*m*. The axon is modeled as a series of cylindrical segments, each 19.9 µ*m* long, with a total of 100 segments. Within a given simulation, all axonal segments share a fixed diameter *d*. Across simulations, the diameter is varied in the range 0.2 µ*m ≤ d ≤* 5 µ*m* in increments of 0.2 µ*m*.

Geometry The presynaptic neuron”s soma is positioned vertically along the *z*-axis, extending from *z* = 0 µ*m* to *z* = 10 µ*m*. The axon originates at the distal end of the soma and extends further along the *z*-axis up to *z* = 2000 µ*m*. All 100 axonal segments are aligned along this axis, resulting in a straight axon oriented in the *z*-direction. The postsynaptic neuron is positioned identically but shifted along the *z*-axis, with its soma extending from *z* = 1000 *µm* to *z* = 1010 *µm*. Its axon originates at *z* = 1010 µ*m* and extends to *z* = 3000 µ*m*.

Membrane potential All compartments were assigned passive and active membrane properties. The specific axial resistivity was set to *r*_*a*_ = 100 Ω *·* cm and the specific membrane capacitance to *c*_*m*_ = 1 µF*/*cm^2^. For a cylindrical segment *n* of length *L* and diameter *d*, with membrane potential *V*_*n*_ and axial neighbors *V*_*n−*1_ and *V*_*n*+1_, the membrane potential evolves according to the cable equation [54]:

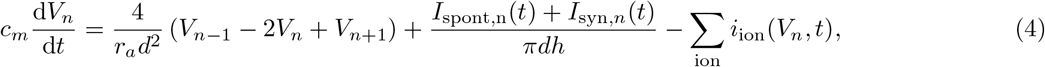

where *i*_ion_ denotes the ionic current density, *I*_spont_ is a current, applied to neurons to elicit spontaneous activity and *I*_syn_ is a synaptic current.

Ionic current densities Ionic current densities are modeled using Hodgkin-Huxley formalism with expressions for the rate constants well suited for cortical pyramidal neurons as described previously [55, 33]. The model includes fast sodium (*i*_*Na*_), delayed-rectifier potassium (*i*_*K*_), and leak (*i*_*L*_) currents, defined as

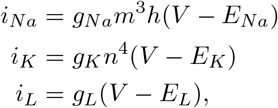

with values for reversal potentials *E* and maximal conductances *g* taken from Traub et al. [55] Gating variables *x* ∈ {*m, h, n*} evolve according to:

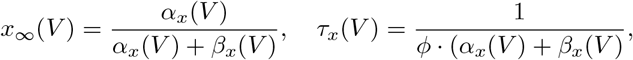

where 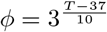 is a temperature-dependent scaling parameter.

Rate functions for each gating variable follow equations from Doorn et al:[33]

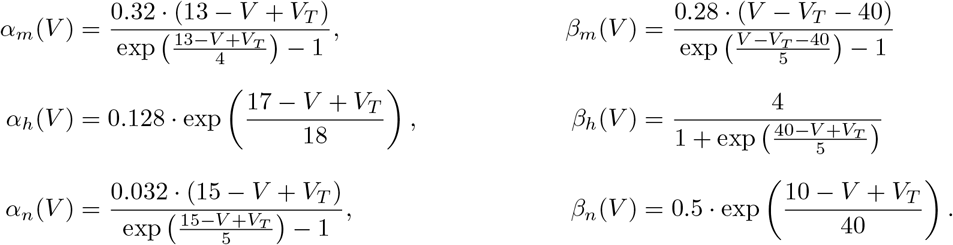

Here, *V*_*T*_ serves as a global voltage offset, uniformly shifting the voltage dependence of all rate functions. In Doorn et al [33], Nernst potentials and a global voltage offset were adjusted to match the average resting membrane potential, spike threshold, and action potential amplitude of NGN2 neurons observed experimentally, and we adopted the same values in our simulations.

#### Synaptic currents

External and synaptic currents are assumed to be inputs from conductance-based chemical synapses. Their currents can be modeled analogously to how ion channels are modeled. For synaptic input, we have

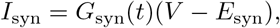

and an analogous equation for external input. The synaptic conductance *G*_syn_ is measured in millisiemens (*mS*). To model the activation of postsynaptic neurotransmitter-gated ion channels, we use a *β*-function profile that captures the rise and decay dynamics of synaptic input following activation. Assuming the synapse is activated at *t* = 0, the conductance evolves as:

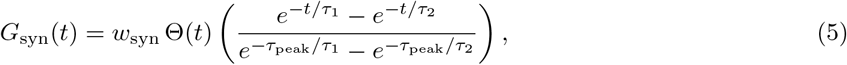

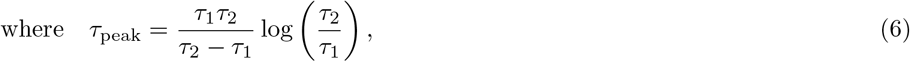

and Θ(*t*) is the Heaviside step function, defined as Θ(*t*) = 1 for *t* ≥ 0 and Θ(*t*) = 0 otherwise. Here *τ*_2_ is the rise and *τ*_1_ is the decay time constants of the synapse.

The inherent stochasticity of synaptic transmission observed in biological neurons, arising from probabilistic vesicle release, fluctuating neurotransmitter concentrations, and variability in postsynaptic receptor activation, was accounted for by sampling the synaptic weight *w*_syn_ (i.e., the peak synaptic conductance) at each presynaptic activation from a truncated normal distribution 𝒯 with specified mean µ and standard deviation *σ*.

To elicit spontaneous activity in each neuron, the initial axon segment receives an excitatory synaptic input *I*_spont_ with reversal potential *E*_spont_ = 0 mV, and synaptic time constants *τ*_1_ = 0.2 ms and *τ*_2_ = 0.4 ms. The synaptic weight for a neuron with axon diameter *d* is sampled from a truncated normal distribution 𝒯 (µ *· d, σ · d*), with µ = 0.02 µS*/*µm and *σ* = 0.02 µS*/*µm. Spike times are drawn from a Poisson process with rate *λ*.

The synaptic weight was scaled with compartment diameter to ensure that the synaptic efficacy is independent of compartment size. Without this adjustment, a fixed conductance input would produce smaller voltage changes in larger compartments due to their increased membrane capacitance and leak conductance. Since the compartment length is fixed across simulations, scaling with diameter alone is sufficient to normalize synaptic impact. The synaptic connection between the two neurons is also modeled using the same formalism, i.e. using Eq. 5, and drawing synaptic weights from a normal distribution with specified mean and standard deviation. *I*_syn_ was further split into AMPA component with fast time constants, and NMDA with slow time constants listed in Tab. 1. The rise and fall timescales for the AMPA component of the synapse were taken to be almost instantaneous, whereas for the NMDA component, these values were taken from [56]. The reversal potential value for both components was also taken from literature [56]. The synapse is located at (0, 0, *z*_syn_) and connects two compartments of the postsynaptic neuron and the postsynaptic neuron, closest to that coordinate.

#### Extracellular electrodes

Two recording electrodes placed in the extracellular space at (0, 0, 60)µ*m* and (0, 0, 1060)µ*m* are modeled to study the extracellular electric fields generated by neuronal activity. The extracellular medium is assumed to be homogeneous and isotropic, with a conductivity of 0.3 S/m. The potential at each recording point is computed using the line-source approximation.

## Model fitting

### Simulation-based inference

Simulation-based inference (SBI) was employed to estimate parameters of a mechanistic neuronal model from observed features of simulated data. Unlike traditional methods that rely on closed-form likelihoods, SBI uses forward simulations to approximate components of Bayes” rule — such as the posterior, likelihood, or likelihood ratio — using a flexible neural network trained on simulation data [57, 36].

To approximate the posterior distribution *p*(*θ* | x), which represents the probability that a given parameter set *θ* produced observed data x, a neural network *q*_*F* (x,*ϕ*)_(*θ*) was trained on simulated (*θ*, x) pairs, where *F* is a parametric family of densities and *ϕ* are its parameters. Because the training data are generated independently of any specific observation x_*o*_, the network learns to map arbitrary inputs to approximate posteriors, enabling *amortized inference* — a setting where inference can be rapidly performed for any new observation within the support of the prior.

Summary features The inference network was trained on summary statistics extracted from simulated voltage traces and corresponding spike times, focusing on biologically meaningful features relevant to synaptic transmission. This approach reduces the dimensionality of the data and targets the aspects most constrained by experimental measurements. As summary statistics, the firing rates and conduction velocities of both neurons were used, as well as the synaptic transmission probability and the coupling lag.

Firing rates of the presynaptic (*f*_pre_) and postsynaptic (*f*_post_) neurons were defined as the number of somatic spikes per neuron divided by the total simulation duration. Conduction velocities (*v*_pre_, *v*_post_) were computed by detecting spike times at two spatially separated sites along each axon, calculating the average time delay between matched spikes, and dividing the physical distance by this latency. For the presynaptic neuron, spikes were recorded between (0, 0, 200) µm and (0, 0, *z*_syn_ *−* 100) µm; for the postsynaptic neuron, between (0, 0, *z*_syn_ + 100) µm and (0, 0, 3000) µm, ensuring that only forward-propagating action potentials were considered.

To compute the synaptic transmission probability, the optimal coupling lag that maximized transfer entropy (as described above) was identified first, then the number of postsynaptic spikes occurring within *±*0.5 ms of that lag were counted and normalized by the total number of presynaptic spikes.

Inference details SBI was implemented using the sbi library [58], employing neural posterior estimation with a mixture density network [59] as the density estimator. The network was trained to minimize the negative log-likelihood of the posterior predictions over *N* simulated samples, 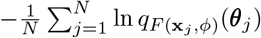, using the Adam optimizer with default hyperparameters [57]. The prior distribution was defined as a uniform distribution over parameter ranges summarized in Table 1. The maximum a posteriori (MAP) estimate was obtained by drawing 10,000 samples from the learned posterior, selecting the 200 with the highest log-posterior values, and performing gradient-based optimization from each to identify the parameter value with the highest posterior density.

The network was trained on a total of 122,000 simulations, each lasting 10 s with a temporal resolution of 0.1 ms. Only simulations for which transfer entropy indicated a significant interaction (*α* = 0.05) were used for inference.

## Results

### Single neuronal pairs can be isolated using microstructures with 10 µm openings

In this work, we introduce a platform for isolating minimal neuronal networks, down to single neuronal pairs, to examine their morphological and electrophysiological characteristics. We achieve this precise isolation by reducing the diameter of the PDMS well to just 10 µm (Fig. 1Ai), a significant modification over previous designs[42, 60, 18, 61] that comprised well diameters ranging from 150 to 400 µm. The approach to neuronal isolation shown in this work presents unique challenges, primarily because the underlying isolation mechanism differs fundamentally from established methods. Unlike systems where cells settle randomly in wells when seeded in suspension [39, 42], or where spheroids can be manually placed using a pipette [18, 61], the system relies on cells that migrate stochastically into wells, predominantly during the initial week in culture.

The efficacy of this microfluidic system depends on two critical factors: ensuring neuronal viability within the microchannels and achieving isolated neuronal pairs that remain functionally independent from surrounding neurons. The migration and survival of neurons into the microchannels required an appropriate matrix. While previous approaches [39, 60, 18, 42] relied on PDL and/or laminin coating to support cell growth, with these we observed low cell numbers in 10 µm wells and poor viability of isolated cells (Figure S1A and B). Higher abundance was achieved when microchannels were filled with matrigel (Fig. 1Bi). Since matrigel was used at a concentration of 3 mg/mL which is highly viscous, we optimized the hydrophilicity of the microchannel surfaces to effectively fill the microstructures. This was accomplished by exposing the substrate and the bottom of the PDMS microstructure (*i.e*., the microchannels) to oxygen plasma prior to mounting the PDMS to the substrate, followed by immediate filling of the channels with diluted, uncrosslinked matrigel. Cells that landed on the top PDMS surface coated with matrigel residuals (Figure 1Bii) gradually migrated downward into the wells (Figure 1Biii). This migration process is documented in the supplementary video, which captures a time-lapse sequence between DIV 0 and DIV 1, immediately after seeding. The time-lapse revealed that over time, increasing numbers of cells migrated into the microwells (Figure 1Ci). The timelapse recordings show that while some neurons established positions in the wells and began extending neurites, others actively navigated along the microchannels (Figure 1Cii). After approximately one week *in vitro*, most microchannels were filled with cells (Figure 1D), resulting in tens to over a hundred potential synaptic pairs in parallel (45% of microchannels contained fewer than two cells, 37% contained exactly two, and 18% contained more than two on DIV 7; N=322).

The irreversible bonding of PDMS microstructures assured the isolation of neurons and prevented neurons from growing in the neighboring microchannels. Plasma treatment following matrigel channel filling selectively degraded the coating on the top PDMS surface as the plasma cannot effectively penetrate the matrigel-filled microstructures. The resulting suboptimal extracellular matrix conditions on top of the PDMS surface induce aggregation and eventual detachment of the remaining cells (Figure 1Biii), thereby preventing unwanted cross-connectivity between the otherwise isolated channels. The comparison of the top of the PDMS microstructure with and without plasma treatment post-matrigel coating is shown in Figure S2. The combination of optimized matrix conditions and selective surface treatment provides a methodology for isolating neuronal pairs that remain viable and functionally independent from surrounding neurons.

### Structural motifs enforce axonal directionality in pre- and postsynaptic neuronal pairs

We aimed to further optimize the system by introducing directionality, thereby ensuring well-defined pre- and postsynaptic neurons and their locations. This directional control is important for creating targeted networks or co-cultures with specific neuronal populations, such as distinct subtypes of cortical neurons or strategic placement of disease-model neurons at predetermined locations within the network.

We designed and tested three main microstructure configurations to establish pre- and postsynaptic neuronal connections (Fig. 2). All proposed designs are compatible with HD-CMOS MEAs to record neuronal signals with high spatial and temporal resolution (Figure 2Ai, Bi, Ci. Since the electrode size (12.5 µm) is comparable with the microchannel width (10 µm) in all designs, each electrode ideally belongs to only one microchannel, therefore narrowing down the variability of neuronal signals it collects.

**Fig. 2:**
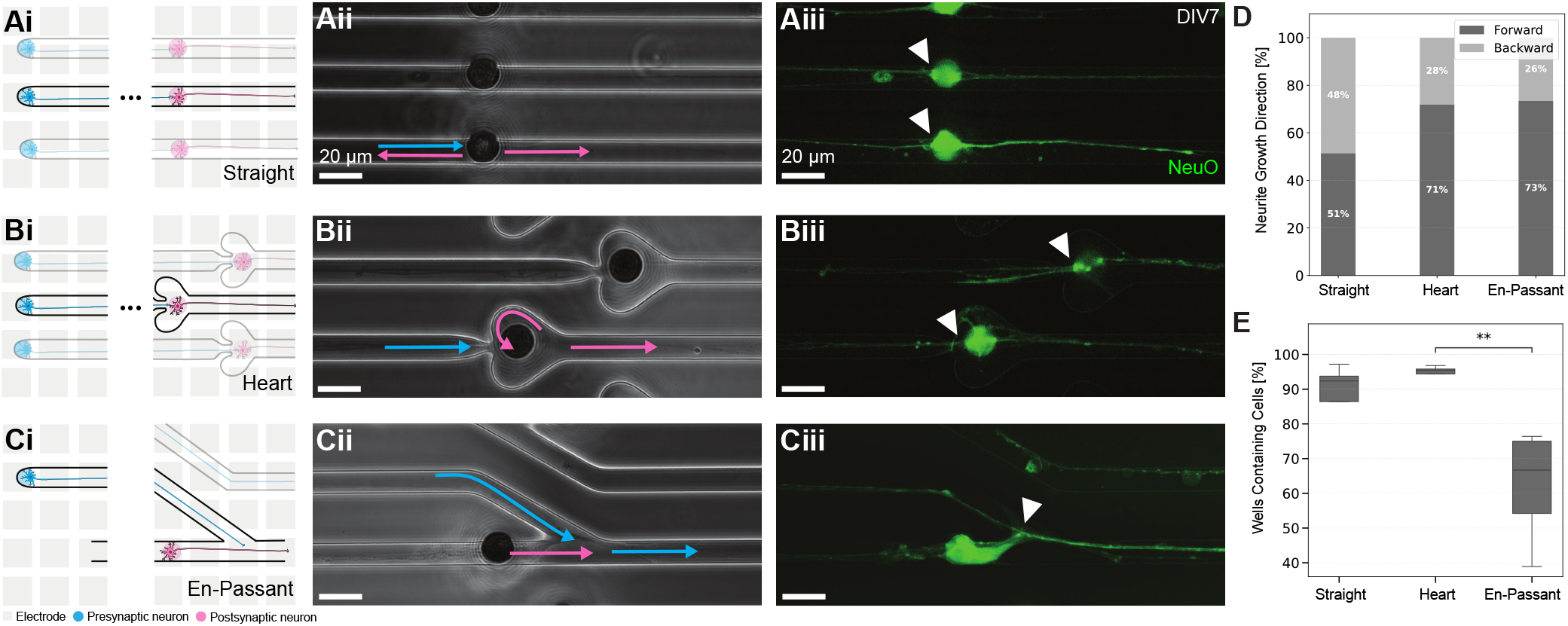
Microstructure designs for directional neuronal growth. A) The straight microstructure design provides a baseline without enforced directionality. Ai) Schematic representation shows a simple channel connecting presy naptic (blue) and post-synaptic (pink) neuronal compartments. Aii) Phase contrast image reveals the fabricated straight microstructure design. Aiii) Fluorescence image demonstrates NeuO-labeled neurons extending processes through the channel, with cell-to-cell contacts visible (white arrowhead). B) The heart-shaped microstructure in corporates axon guidance principles. Bi) Schematic illustration depicts the heart-shaped feature designed to prevent backward axonal growth. Bii) Phase contrast image shows the fabricated heart-shaped microstructure with guid ance features for the postsynaptic neurons highlighted (pink arrow). Biii) Fluorescence imaging confirms neuronal growth through the structure with presynaptic neurites extending to the postsynaptic neuron (white arrowheads). C) The en-passant microstructure design facilitates axon-dendritic connections. Ci) Schematic representation shows the angled channel design for axon-dendritic en-passant synapses. Cii) Phase contrast micrograph demonstrates the fabricated en-passant structure with highlighted directional cues. Blue and pink arrows indicate desired growth directions for pre-respectively postsynaptic neuron). Ciii) Fluorescence imaging confirms neuronal growth through the angled channels with the postsynaptic cell body in contact with the presynaptic neurites (white arrowhead). D) Quantification of neurite growth direction across the three designs shows: straight (51% forward, 49% backward, N=175), heart (72% forward, 28% backward, N=245), and en-passant (73% forward, 27% backward, N=94). E) Quantification of channel occupancy (percentage of wells with cells) reveals a significant difference (*p *<* 0.05) between designs, with en-passant showing lower occupancy compared to heart and straight designs. Error bar represents 95% confidence interval. Scale bars: 20 µm.

The first design (Figure 2A) featured a straight microstructure with no enforced directionality other than the well locations. The second design (Figure 2B) incorporated a structure based on axon guidance principles, which states that axons tend to follow edges, but will not follow the edge around sharp corners [62, 14, 63, 64]. Due to space constraints, we implemented a minimal design where a heart structure with a narrow entering region reduces the probability of the postsynaptic axons to grow back into the presynaptic channel (indicated by pink arrows in Figure 2Bii). The third design was inspired by the axo-dendritic “en-passant” synapses described previously [65], where edge guidance and differences in axonal and dendritic lengths create unidirectional connections from pre-to postsynaptic neurons with “en-passant” synapses. In this design, directionality of both the pre- and postsynaptic neurons is controlled by introducing a sharp turn in the junction of the channels (indicated by pink and blue arrows in Figure 2 Biii). The desirable contact points between the presynaptic axon and the postsynaptic cell are indicated by white arrows in Figures 2Ci-Ciii for each design.

We quantified neurite growth direction using a combination of Fiji and manual annotation to assess the success of the three designs in enforcing directionality in the formation of synapses (Figure 2D). The straight design exhibited no preferential directionality, with nearly equal forward (51 %) and backward (49 %) growth (N = 175). The close to 50:50 bidirectional growth pattern confirms that without structural guidance, neuronal processes extend stochastically and equally in both directions. In contrast, both the heart design (N = 245) and en-passant design (N = 94) demonstrated significant directional preference, with comparable 72 % and 73 % forward growth. While robust, the directionality is lower than the directionalities above 90 % reported in previous studies [65, 14, 66]. This difference might be attributed to our simplified designs, which prioritized spatial efficiency and ease of fabrication.

Although the percentage of channels containing cells was similar across all designs, we observed a significant increase in cell occupancy within the heart and straight designs compared to the en-passant configuration (Figure 2E). We hypothesize that this difference results from subtle variations in design. The straight and heart designs feature two rows of wells, while the en-passant configuration consists of three rows (see Fig. S4-S6). Since the wells are more distributed, the probability of cell that landed on the top surface migrating to a well is lower.

Despite these minor differences, the comparable channel occupancy across all designs indicates that our structural modifications did not significantly affect neuronal viability or the accessibility of the channels. We present a wide spectrum of channel designs, from standard microchannels down to nanochannels, which enable precise control over axodendritic synaptic connections, in Figures S4, S5, S6, S7, and S8. These designs support diverse experimental paradigms from basic synaptic morphology studies in monocultures to complex co- or tri-culture investigations.

### Isolated neurons are active over a long period in culture, allowing detailed characterization

Neurons typically develop and function within complex networks, rarely surviving in isolation under standard culture conditions due to their dependence on trophic support and network activity [67, 68]. Therefore, a critical concern for the isolated neuron approach was whether these cells would remain viable and functionally active over extended periods.

We showed that neurons constrained in our microenvironment not only survive but maintain robust electrical activity for several months. Placing PDMS microstructures on top of HD-MEAs enabled comprehensive electrophysiological characterization of the cultures. The physical isolation of neurons within microchannels ensured that we were consistently recording signals from the same neurons throughout the experiment. The spike sorting approach we employed automatically detected and assigned extracellularly recorded action potentials to individual neurons (Figure 3Bi and Bii). Waveform shapes varied from one neuron to another but also changed along the microchannel for the same neuron, which could be attributed to recording from different cellular compartments (from dendrites, through somas to axons) [34], varying positions of the recorded cell parts with respect to the electrodes, or the relative position of the electrodes with respect to the PDMS microstructure [69]. Figures 3Ci and 3Cii illustrate representative waveform metrics extracted from our recordings. Additional extracted waveform metrics are available in Figure S9. Notably, the majority of recorded action potential amplitudes exceeded 100 µV, whereas these values are only achievable over multiple averages in random cultures [34], highlighting the significant advantage of our PDMS microstructures in improving the signal-to-noise ratio.

**Fig. 3:**
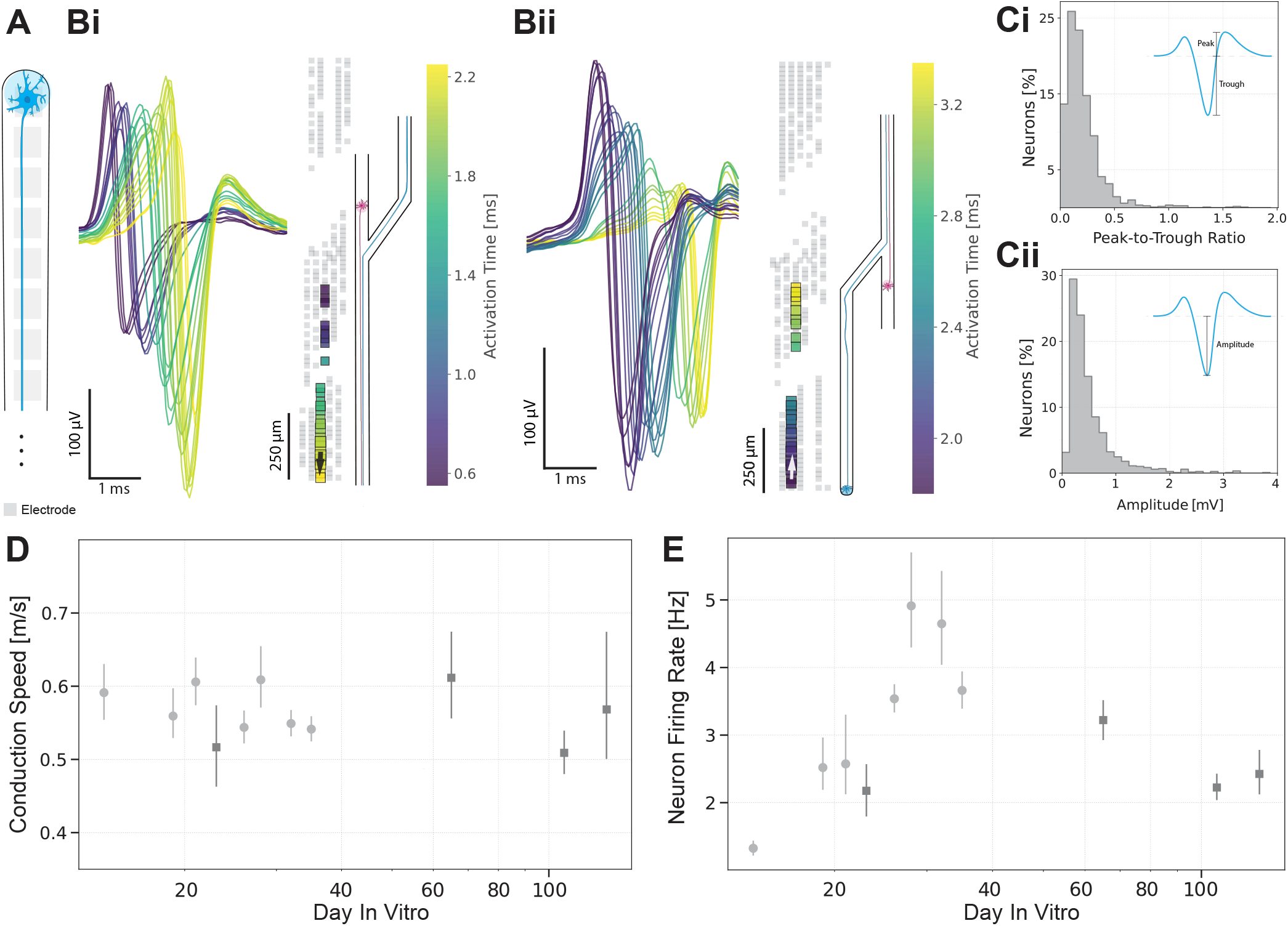
A) Schematic of a single neuron in a microchannel. B) Activity of single neurons in microchannels. Bi) One example neuron recorded at different microchannels clustered using the Spikeinterface framework [43]. Templates from twenty five electrodes belonging to the same channel are plotted in time. Time progression of spikes is color-coded. Black arrow shows the direction of action potential propagation within the microchannel. The schematic shows the assumed part of the network that the neuron belongs to. Bii) An example of another neuron on the same chip with a different template progression. C) Example metrics extracted per unit/neuron at DIV 65 (N = 1432), showing distributions of Ci) peak-to-trough ratios and Cii) amplitudes. D) Evolution of conduction speed with culture age. One set of chips (N = 4) is used to assess the behavior during the first five weeks (light gray circles) and another (N = 1) over longer time in culture (up to 129 days, dark gray squares). E) Evolution of the mean neuron firing rate of the neurons in D) over 129 days. Error bars in D) and E) represent 95% confidence interval.

Neurons exhibited sufficient activity for reliable calculation of action potential propagation (Figure 3 D) and firing rate (Figure 3 E) from the second week (DIV 14) onward. We observed a relatively constant conduction speed between DIV 14 and DIV 129 which, given the established relationship between conduction velocity and unmyelinated axon diameter [70, 71], indicated that axonal morphological properties and ion channel distribution along the axons were established early in development. In contrast, the firing rate showed a steady increase until DIV 28, reflecting ongoing functional maturation through the formation and strengthening of synaptic connections. After complete maturation, the firing rate stabilized between 2 and 3 Hz and remained relatively constant throughout the culture”s lifespan. The minor periodic fluctuations observed in both conduction speed and firing rate over weeks *in vitro* likely resulted from environmental factors such as temperature fluctuations or changes in medium composition due to evaporation and cellular processes, rather than developmental changes in the neurons themselves.

These findings are consistent with previous reports on NGN2-induced neurons [72, 67, 61], some of which also describe conduction velocities around 0.5 m/s and spontaneous firing rate stabilization after four weeks in culture.

Importantly, we demonstrated that neurons remained viable for over 120 days, which is longer than the durations reported in most studies using NGN2 neurons without additional support from glial co-culture or specialized media [67, 73, 74, 68]. This, and the stability of measurements, confirmed the effectiveness of PDMS isolation for long-term neuronal recordings.

### Isolated neuronal pairs form functional synapses

Having demonstrated that neurons mature and maintain viability for extended periods within the microstructures, we next investigated whether these isolated neuronal pairs in microchannels establish functional synaptic connections. Previous computational, *ex vivo* and *in vitro* work has demonstrated that multiple synapses and simultaneous incoming stimuli are typically required to elicit action potentials in postsynaptic neurons [23, 75, 76, 77]. Therefore, it was crucial to verify both the physical presence of synapses and their functional significance in the isolated two-neuron system.

We identified putative synaptic pairs by testing whether the past activity of a presynaptic neuron significantly reduces uncertainty about the current state of a postsynaptic neuron, beyond what can be inferred from its own past. This was quantified using transfer entropy — a model-free measure of directed information flow between time series [78, 79] (see Methods). This approach has already been applied to large-scale neural data [80, 81].

To validate the functional connections identified by transfer entropy, an artificial neural network was trained to forecast postsynaptic spike probabilities based on preceding pre- and postsynaptic activity. The results of this analysis further support the transfer entropy-based identification of synaptic pairs: for neuron pairs classified as coupled by transfer entropy, both traces significantly contributed to prediction accuracy (see Supplementary Material S15). While transfer entropy captures directed information flow, it does not by itself confirm causal synaptic transmission [82]. We therefore complemented the functional analysis with immunocytochemistry, which revealed colocalization of synapsin and PSD95 puncta—markers of pre- and postsynaptic proteins—across all microstructure designs, confirming the physical presence of synaptic contacts (Figure 4A).

**Fig. 4:**
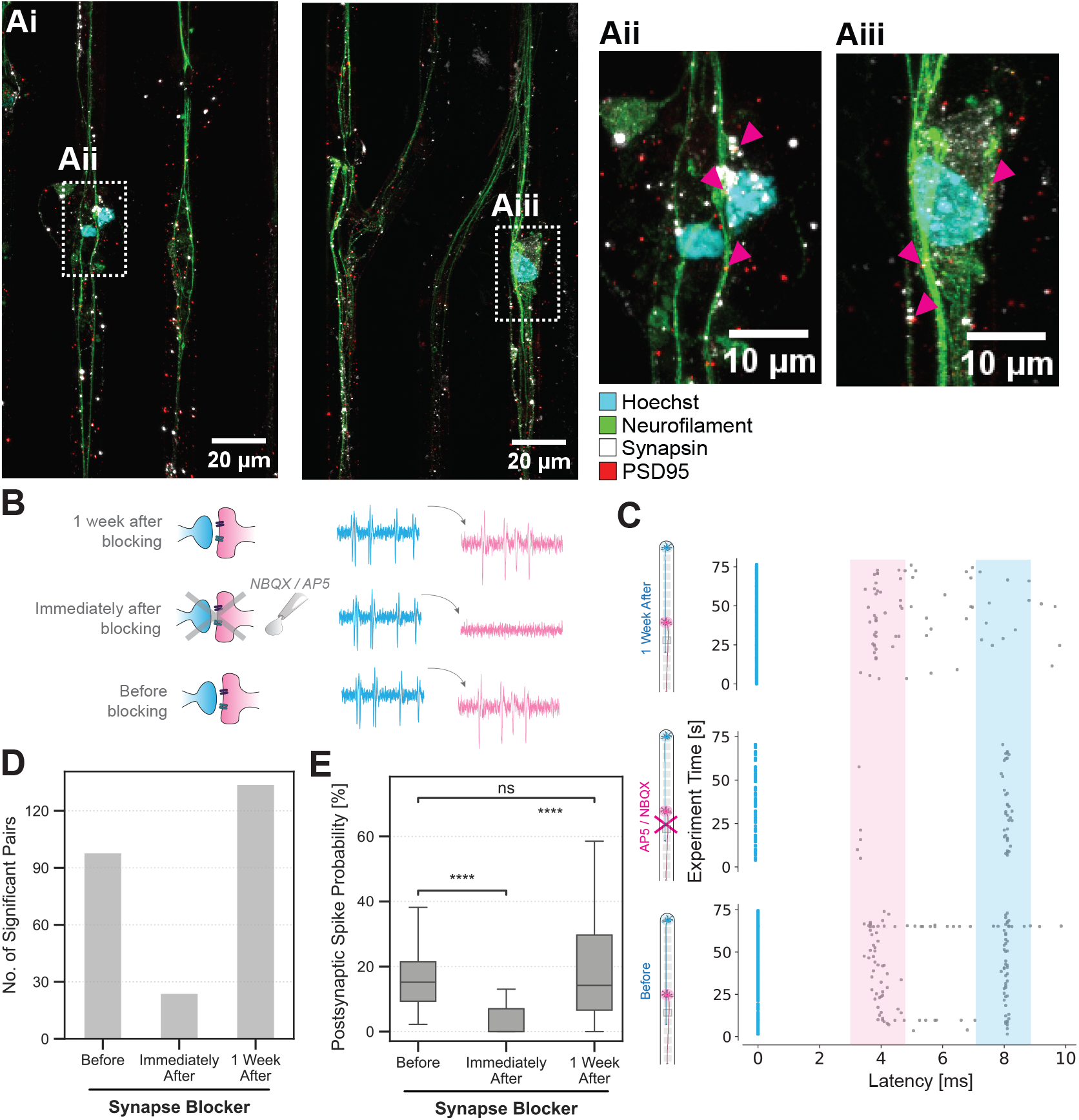
Validation of synaptic connectivity in isolated neuronal pairs in the and en-passant microchannels.A) Immunocytochemical evidence of synaptic connections. (i) Fluorescent microscopy images showing the colocalization of synaptic markers. (ii) A higher magnification inset provides detailed visualization of the synaptic structures. B) Schematic of the experiment. NBQX and AP5 were added as NMDA and AMPA receptor antagonists on DIV 100 and the activity was recorded before the application of synaptic blockers, immediately after and one week after washout. C) Representative spontaneous spike time-triggered latency plot shows spikes before, in the presence of synaptic blockers and one week (following washout) after the application of synaptic blockers, demonstrating the change in synaptic transmission properties. D) Bar graph showing the number of significant transfer entropy links before, after synaptic blocker application, and following washout for one chip. E Box plots quantifying the synaptic transmission probability across the three conditions, with statistical significance indicated (****p*<*0.0001).

To examine the functionality of putative synapses in the system, we performed a pharmacological intervention on DIV 100 using AMPA and NMDA receptor antagonists (NBQX and AP5, respectively) to inhibit glutamatergic synaptic transmission (Figure 4B). We recorded and compared spontaneous neuronal activity across three time points: before antagonist application, in the presence of the antagonists immediately after application, and one week following washout. Figure 4C) shows a spontaneous-spike-time-triggered latency plot illustrating the relationship between one source-target (*i.e*., pre-and postsynaptic neuron) electrode pair. To eliminate delays from axonal propagation, we chose the same electrode as both source and target, as it captured signals from two different neurons within the microchannel. Latency plot revealed two distinct activity bands at approximately 4 ms and 8 ms timepoint. The 4 ms band (highlighted in pink) appears wide and relatively sparse. In contrast, the 8 ms band (highlighted in blue) is denser, suggesting it represents activity from the same presynaptic neuron that is used as a trigger at 0 ms. This interpretation is substantiated by the differential response to receptor antagonists: immediately following antagonist application, the 4 ms band is markedly diminished, indicating successful blockage of synaptic transmission, while the 8 ms band persists, albeit with reduced density—reflecting an overall decrease in neuronal activity following pharmacological intervention.

We focused on two metrics to quantify these functional changes: the number of significant pre- and postsynaptic neuronal pairs, defined as those with transfer entropy significant at the 0.05 level, across the three time points; and the probability of postsynaptic spike occurrence following presynaptic spikes. The number of significant pairs shows a four-fold decrease immediately upon antagonist addition and a six-fold recovery one week after washout (Figure 4D). Similarly, the administration of synaptic blockers resulted in a significant reduction in the postsynaptic spike probability (Figure 4E) as well as a decrease in pairwise transfer entropy values (shown in Figure S10). Significant differences were observed between pre-blocker and blocked conditions as well as between blocked and washout conditions. Importantly, no significant difference was found between pre-blocker and one-week-after washout conditions, indicating complete recovery of synaptic function after blocker removal. The reversible disruption of postsynaptic spike probability and transfer entropy by synaptic blockers and subsequent recovery after washout demonstrate that neuronal pairs isolated in microchannels form functional synaptic connections, despite their minimal configuration.

Putative synaptic transmission can also be confirmed by visual inspection on a case-by-case basis. Spike sorting of the recordings enabled to distinguish action potentials from neurons growing in close proximity (Figure 5A). Figure 5B illustrates spike propagation and putative synaptic transmission in a representative microchannel. The recordings demonstrate action potential propagation along the microchannel, with the initial spike traveling from the top electrode (negative deflection at 1.75 ms) and continuing down the channel (positive component at 2.75 ms). A secondary action potential emerges at 4.75 ms in the bottom half of the microchannel, exhibiting propagation and a 3 ms synaptic delay that indicates postsynaptic initiation rather than passive propagation of the original signal.

**Fig. 5:**
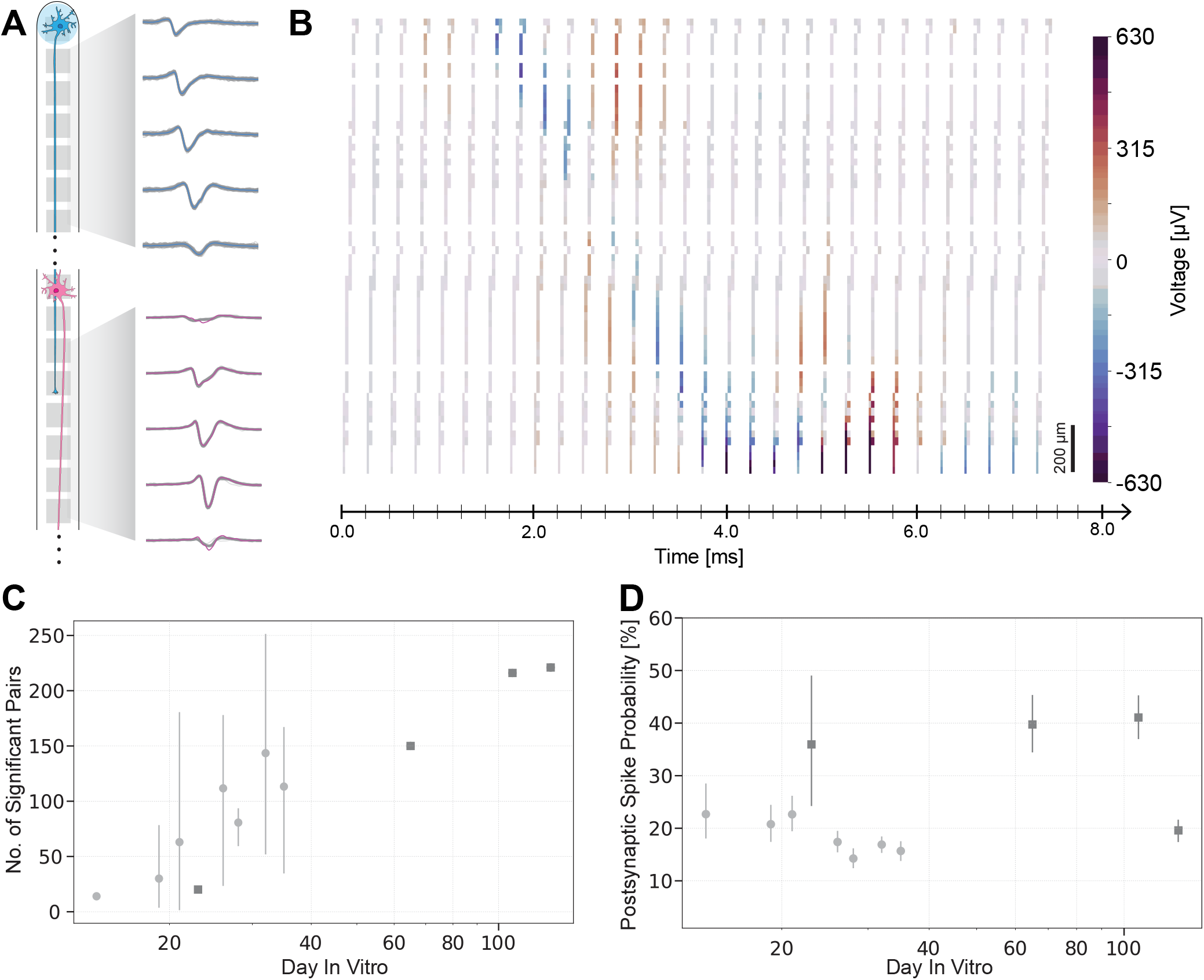
Synaptic connectivity dynamics in isolated neuronal pairs. A) An example of processed electrophysiological recordings from neurons in microchannels showing spike-sorted traces from both pre-synaptic (blue, top) and post-synaptic (pink, bottom) neurons with the propagation of the traces along the channel by displaying 5 equally spaced electrodes for each neuron. B) Video frames showing the temporal relationship between pre- and post-synaptic activity with color-coded amplitude scale in microvolts (µV). Scale bar: 200 µm. C) Number of significant pairs found by transfer entropy analysis for one set of chips (N = 4) early in culture (first five weeks, light gray circles) and for another (N = 1) over longer time in culture (up to 129 days, dark gray squares). D) Postsynaptic spike probability of the neurons in C) early in culture (first five weeks, light grey circles) and over longer time in culture (up to 129 days, dark gray squares). Error bars in C) and D) represent 95% confidence interval.

We systematically analyzed different synaptic metrics over time to quantify the long-term dynamics of functional connectivity. The number of functional pairs reveals a clear developmental trajectory during the first five weeks in culture (Figure 5C). While few significant connections are detected in the earliest recordings (DIV 15-20), the number increases substantially during the first month in culture. The number of detected pre- and postsynaptic pairs further increases after DIV 80, indicating the ongoing network formation and enhancement of functional connectivity throughout the culture period. One hypothesis is that putative remaining neurons on top of the PDMS surface continue to migrate and project axons inside the microchannels. Another hypothesis is related to the fact that individual NGN2 neurons can develop multiple axons even after 4-6 weeks in culture [83, 11]. Similarly, postsynaptic spike probability varies between 15 - 35 % within the first 20 days, increases to 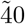 % beyond DIV 60 as networks mature, and notably declines at DIV 129, potentially indicating culture senescence (Figure 5 D).

This observation is particularly interesting when contrasted with the number of significant connections, which remains high at this time point, and the firing rate (Figure 3E) suggesting that while the quantity of neuronal pairs persists, the efficacy of their synaptic transmission decreases. This implication aligns with the literature that states that neurons suffer synaptic dysfunction and axonal connectivity loss prior to somatic cell death [84, 85].

### Simulation-based inference finds the biophysical parameter space from microelectrode-array data of neuronal pairs

We complemented the experimental platform with a biophysical Hodgkin-Huxley (HH) model [86] to further leverage the rich electrophysiological data from the experiments and investigate the mechanisms of synaptic transmission.

The absence of complex neuronal networks in our *in vitro* system allowed for the use of a minimal computational model capable of capturing the characteristics of signal propagation and transmission. Specifically, we modeled the network consisting of two neurons using a simple multicompartmental formalism, with each neuron having two compartments: a ball representing a soma and a stick representing an axon. Passive and active properties suitable for human cortical pyramidal neurons were adapted from existing models [33, 55]. Pre- and post-synaptic neurons were connected by excitatory synapses to study the synaptic transmission between the neurons (see Methods for details). The model embodied Occam”s razor — complex enough to study synaptic transmission in this experimental system, yet simplified by omitting dendritic structure and multiple synapse locations.

For model fitting, we used simulation-based inference (SBI) [87, 36, 37, 88]. Instead of direct likelihood calculation that would require the ability to integrate over all potential paths through the simulator code, in SBI, forward simulations are used to train a model (typically, a neural network) to approximate parts of Bayes” rule. Here, we used SBI to approximate the posterior distribution (Figure 6). This approach was found to be particularly useful in neuroscience, since it is possible to infer parameters in highly complex models without closed-form likelihoods [87]. We extracted six key summary statistics (Fig. 6A) from the electrophysiological recordings from neuronal pairs on MEAs (Fig. 6B), including: firing rates, conduction speeds, postsynaptic spike probability and synaptic delays. These experimental statistics defined the loss function for training a neural density estimator (NDE), ensuring the NDE identified the most likely probability distribution consistent with the provided data. To train the model, we generated a simulated dataset by sampling from uniform prior distributions over biophysically relevant parameters (Fig. 6C) and feeding these into the mechanistic Hodgkin-Huxley model (Fig. 6D). The NDE (Fig. 6E) was then trained on this simulated data to approximate the probability distribution of model parameters given specific summary statistics. Using the model trained this way, we could infer the corresponding model parameters for each recorded neuron pair along with the corresponding posterior distribution of model parameters (Fig. 6F) as a comprehensive view of the parameter landscape that is consistent with observed data.

**Fig. 6:**
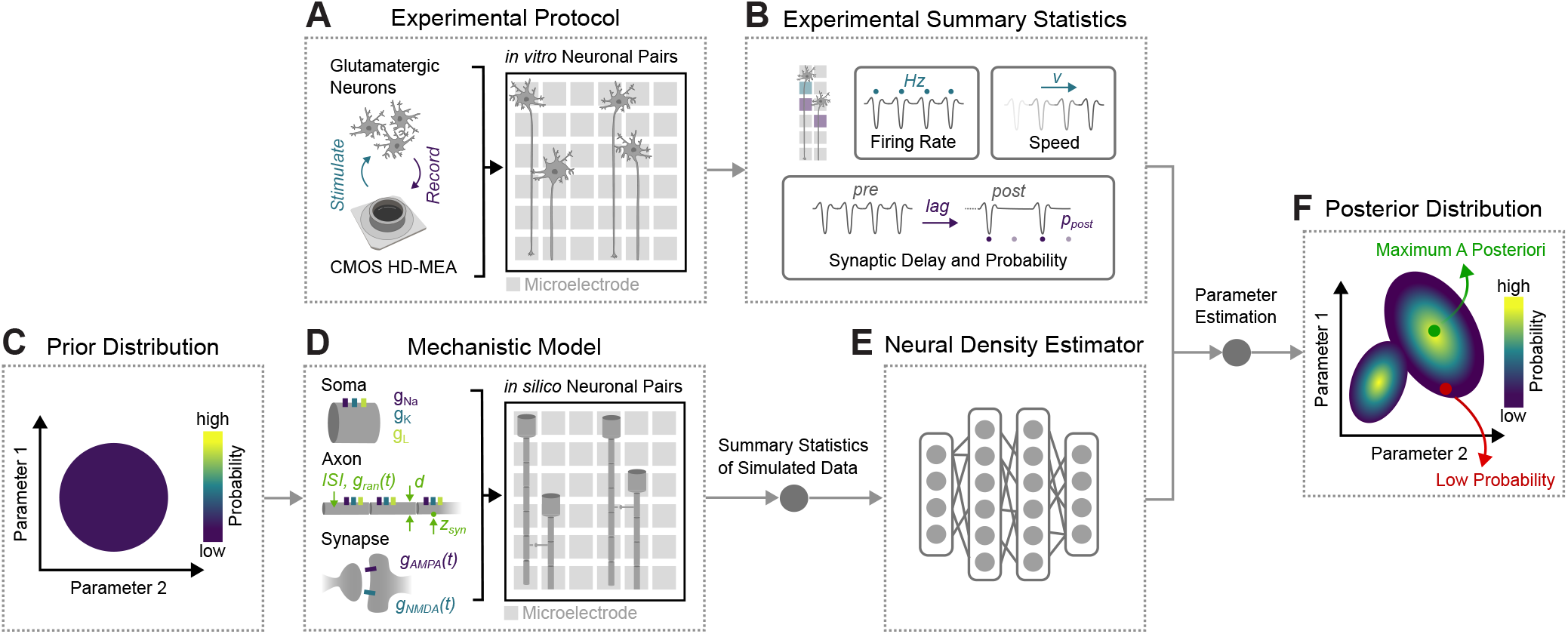
Schematic overview of the method (adapted from Gonçalves et al. [87]) A)First, NGN2 neurons are cultured on CMOS high-density microelectrode arrays (HD-MEAs) in PDMS microstructures, enabling stimulation and recording from *in vitro* neuronal pairs. B) Experimental summary statistics were extracted from the spontaneous activity recordings. C) 100,000 parameter configurations were drawn from a uniform prior distribution of model parameters. D) Using a mechanistic model of isolated neuronal pairs incorporating Hodgkin-Huxley-type neurons, these parameters drove *in silico* neuronal pair simulations. E) A neural density estimator was trained on summary statistics from the simulated data. The trained neural density estimator was used to predict parameter estimations from experimental summary statistics. F) Posterior distribution resulting from parameter estimation with the neural density estimator, highlighting the maximum a posteriori estimate and regions of low probability in the parameter space.

First, we demonstrated that the experimental setup (Figure 7A) can be successfully emulated by a biophysical model (Figure 7B), whose parameters were inferred using SBI. This was first shown for one exemplary neuronal pair (Figure 7Ci) exhibiting characteristic latency patterns between pre- and postsynaptic neurons over time. Simulations using maximum a posteriori (MAP) parameter configurations replicated these experimentally observed activity patterns (Figure 7Cii), while simulations using low-probability parameter sets substantially deviated from experimental data (Figure 7Ciii). Analysis of the full posterior distribution (Figure 7D) shows that most parameters, such as the presynaptic and postsynaptic diameters (*d*_pre_, *d*_post_), average inter-event times of spontaneous neuronal activity (1*/λ*_pre_, 1*/λ*_post_), average AMPA/NMDA synaptic weights (*µ*_AMPA_, *µ*_NMDA_), and synapse loca tion (*z*_syn_)—exhibited narrow posteriors, indicating that the experimental data strongly constrain these parameters and the recorded observables exhibit minimal redundancy or compensation from other parameters. For instance, neuronal diameters primarily determine conduction speed, which is directly captured in the experimental summary statistics. Other parameters displayed broader distributions, particularly the standard deviations of the weight of the AMPA and NMDA receptors (*σ*_AMPA_, *σ*_NMDA_), implying that multiple parameter combinations can reproduce the experimental data or that the experimental summary statistics are weakly sensitive to these parameter variations. To further understand how these parameters affect summary statistics, we analyzed correlations between summary statistics x and model parameters *θ* across all simulations (Figure 7E). The resulting correlation matrix revealed significant associations, with strong correlations confirming expected biophysical relationships such as the tight coupling between axon diameter and conduction speed, and AMPA receptor weight with synaptic probability [89, 90].

**Fig. 7:**
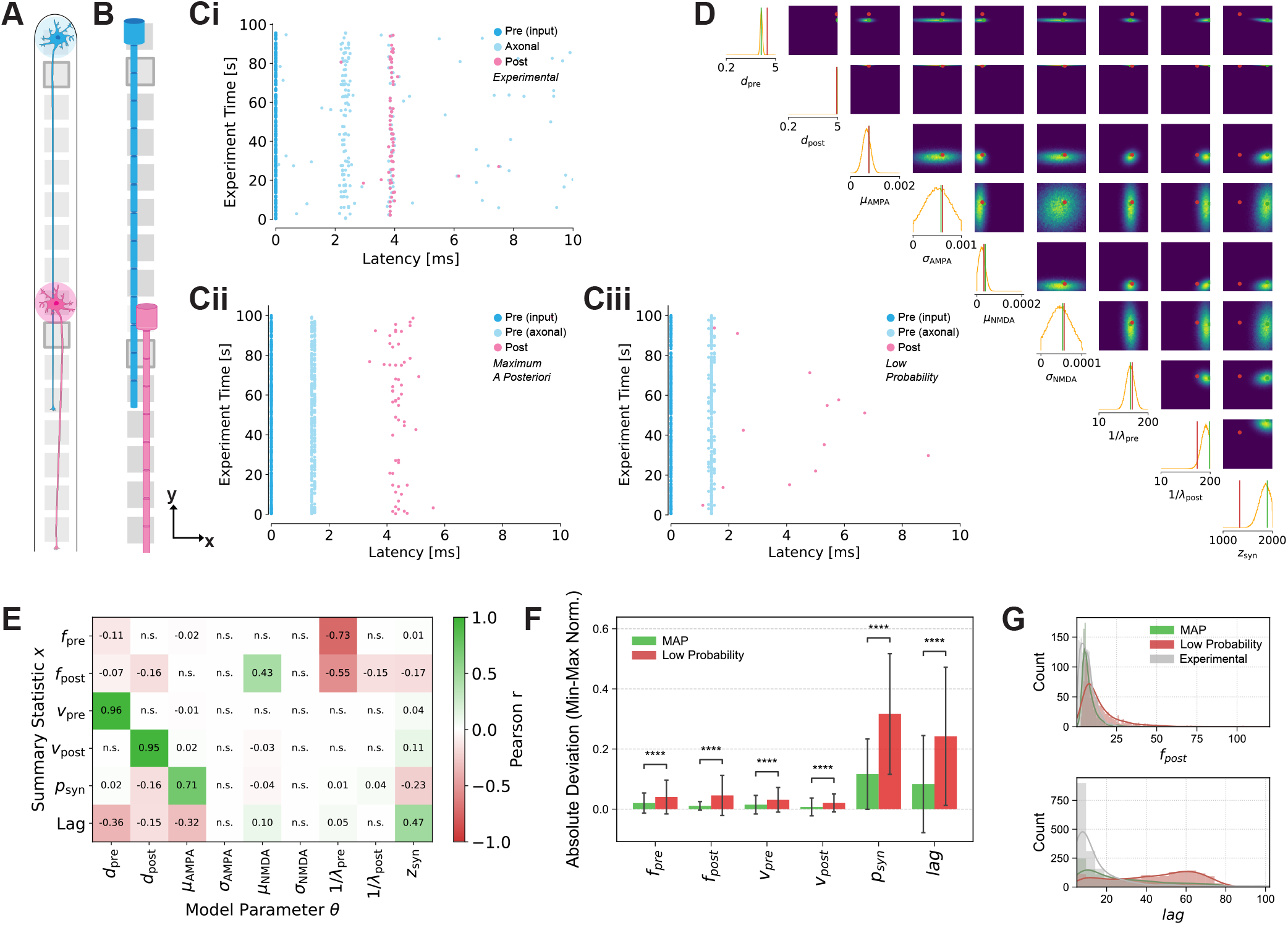
Neuronal pairs parameter inference and validation: A) Pre-synaptic (blue) and post-synaptic (pink) neurons in the experimental setup are modeled using a B) mechanistic neuronal network model with presynaptic (blue) and post-synaptic (pink) ball and stick neurons. C) Comparison of latency plots that were obtained experimentally and with the model. Ci) Experimental data displaying pre-input (dark blue), axonal (light blue), and post-synaptic (pink) spike latencies across the experimental time can be approximated with the mechanistic models using Cii) maximum a posteriori parameter configuration. The maximum a posteriori parameter configuration shows improved alignment between predicted and experimental latencies compared to Ciii) low probability parameter set simulations. D) Inferred posterior of nine model parameters (axon diameter of the pre- and postsynaptic neuron, respectively, mean and standard deviation of AMPA- and NMDA-receptor weight, respectively, intrinsic activity parameter for pre- and postynaptic neuron, respectively) derived from the experimental data shown in Ci). Univariate respectively pairwise marginals are shown on the diagonals and off-diagonal, where maximum probability parameter configuration is shown in green (dot and line) and a low probability parameter configuration example is shown in red. E) Correlation matrix shows dependencies between six summary statistics (firing rate and conduction speed of pre- and postsynaptic neuron, respectively, postsynaptic spike probability and the spike delay) and the nine model parameters mentioned in D. F) Summary statistics derived from model simulations using maximum a posteriori (green) parameter configurations deviate significantly less from summary statistics derived from experimental data compared to statistics derived from model simulations using low probability (red) parameter configurations. Statistical significance indicated (****p*<*0.0001). G) Distribution histograms comparing frequency counts between MAP model (green), Low Probability model (red), and Experimental data (gray) for two example statistics.

**Fig. 8:**
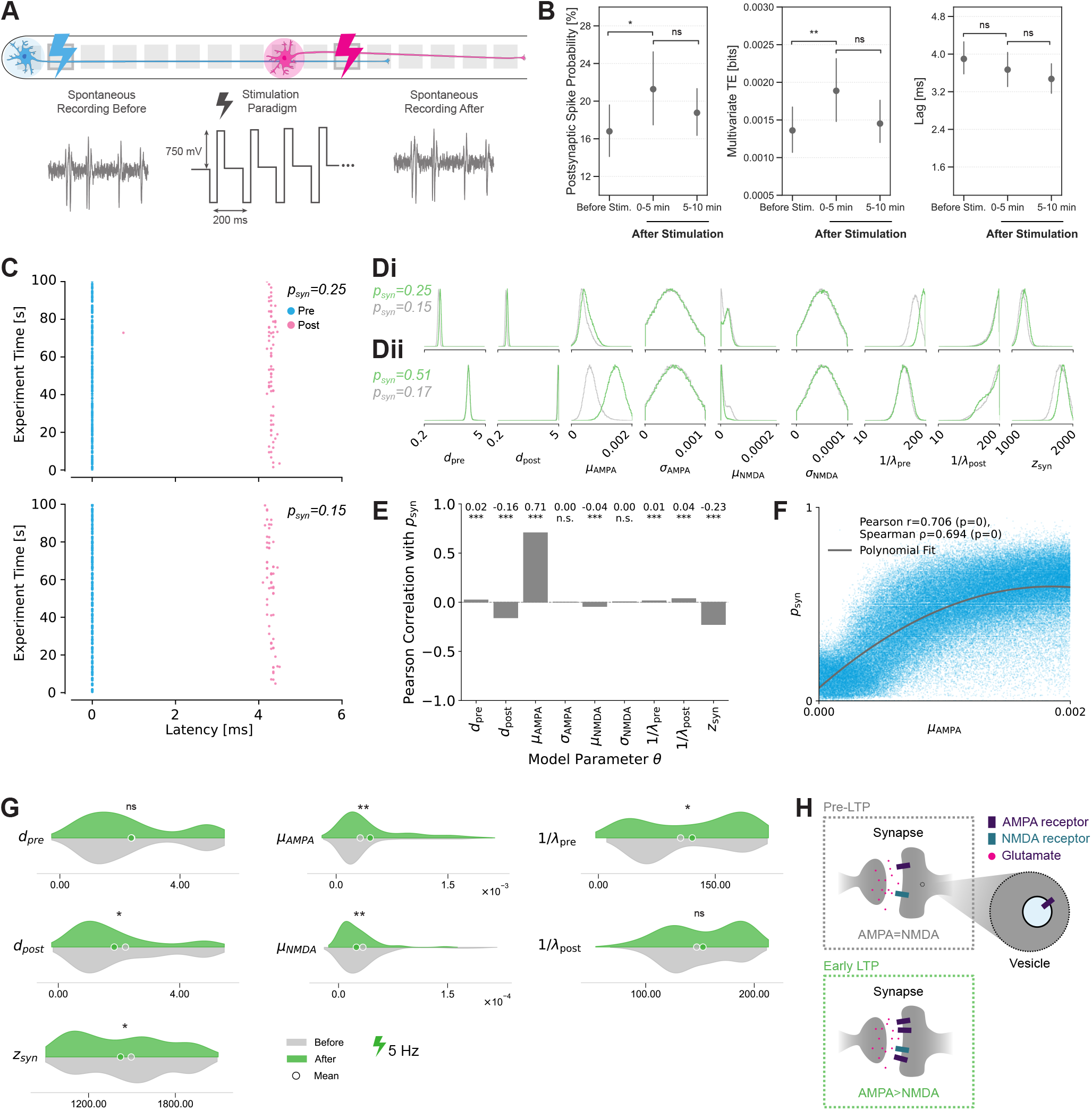
A) Experimental stimulation paradigm consists of 5-second 5 Hz simultaneous stimulation of electrodes that belong to pre- and postsynaptic neurons with 5-second break in-between for 5 minutes. The spontaneous activity is recorded and analyzed before and two times consecutively after stimulation. B) Synaptic metrics quantifying changes upon stimulation. Postsynaptic spike probability and multivariate transfer entropy show significant increases immediately following stimulation, while lag remains statistically unchanged. C) Latency plots of the same neuronal pair before and after stimulation. The postsynaptic spike probability values are 0.15 and 0.25 before and immediately after stimulation, respectively. D) Single sample marginal distribution changes. Di) Distribution changes for an experimental example from C). Dii) Distribution changes for a three-fold change in *p*_*syn*_. E) Pearson correlation between the postsynaptic spike probability and the model parameters computed across prior simulations. F) Postsynaptic spike probability increases with the weight of AMPA in the HH model, reaching saturation at high values of AMPA. G) KDE density plots of the MAP parameter configurations identified in all included neuronal pairs before and after stimulation with 5 Hz. Statistically significant differences between the means are indicated (*p*<*0.05, **p*<*0.01, ***p*<*0.001). H) Schematic of the AMPA and NMDA regulating synapse dynamics and stabilization (adapted from [91]).

Using this framework, we were able to infer likely parameter values for over 1000 experimental pairs at DIV 35. The min-max normalized absolute deviation was clearly below or close to 0.1 for all summary statistics when using MAP parameters for the simulations, demonstrating accurate reproduction of experimental features (Figure 7F), non-normalized values in Figure S11A). In contrast, simulations from low-probability parameter sets showed much larger deviations, particularly for synaptic probability and delay (lag). The smaller absolute deviation for the low probability parameter configuration in conduction speed is consistent with the narrow marginal distributions observed for the pre- and postsynaptic diameter parameters (Figure 7D), which are the main parameters determining conduction speed (Figure 7E). Additionally, we also quantified the information loss when approximating the experimental data with the models using the KL divergence (Figure S11B). The information loss using the MAP parameter configuration was lower or equal to the low probability configuration for all parameters except the presynaptic neuron activity, which again can be explained through the dependency on parameters with narrow marginal distributions.

These quantitative findings were further supported by distribution histograms (Figure 7G) from the same set of simulated and experimental pairs, which illustrate that the MAP model aligns closely with experimental data for both post-synaptic firing frequency and synaptic probability, whereas the low-probability model exhibits marked discrepancies. Other distribution histograms were less distinct (Figure S11C). These results confirm that simulation based inference can robustly identify model parameters that replicate neuronal network activity as measured on

### Neuronal model can explain synaptic potentiation upon stimulation

Previous studies in more complex preparations have demonstrated that synchronous stimulation of pre- and postsy-naptic neurons can induce LTP [7, 10, 5]. We sought to replicate this phenomenon in our simplified *in vitro* system and characterize the observed changes using the SBI approach to investigate the possibility of targeted modulations of synaptic strength.

We implemented a stimulation paradigm comprising three phases: an initial five-minute spontaneous recording to establish baseline activity, followed by simultaneous stimulation of pre- and postsynaptic neurons through the identified source and target electrodes (Figure 8A), and concluding with two sequential five-minute recordings, one to capture immediate post-stimulation effects and another to quantify the potential recovery to pre-stimulation baseline conditions. Both postsynaptic spike probability and multivariate transfer entropy (see Methods) significantly increased after stimulation (Figure 8B), indicating enhanced synaptic efficacy. While there is an observable decrease of both metrics in the second recording after stimulation, the values are not significantly different neither from the ones before nor from the ones immediately after stimulation. This indicates the system slowly returning to basal state, which is also visible in the decreasing postsynaptic spike probability curve over time shown in Fig. S12A). Interestingly, the changes in the transfer entropy lag did not reach statistical significance, potentially because the temporal resolution (1 ms) could be insufficient to capture the potential shifts in the synaptic delay. We additionally analyzed the changes in firing rate and conduction speed within the same recordings for pre- and postsynaptic neurons, respectively (Fig. S12B-C). The postsynaptic neuron firing rates exhibit more pronounced changes, potentially reflecting modifications in synaptic strength. In contrast, conduction speeds remain stable throughout the experimental period. Since conductance of ionic channels is highly sensitive to environmental factors such as temperature fluctuations [92], this relative stability indicates that the observed metric changes result from the effect of stimulation on neurons rather than confounding environmental variables. This suggests that the observed effect is caused by a synaptic potentiation mechanism.

Using the HH model and the previously described SBI approach, we could decompose the changes in postsynaptic spike probability and multivariate transfer entropy into contributions from different synaptic components. Figure 8C shows an example latency plot of a neuronal pair with 10 % increase after stimulation. Even in such cases when the changes in experimental postsynaptic spike probability were relatively small, the corresponding marginal parameter distributions showed informative changes (Figure 8Di). Notably, there was a shift towards higher values for the mean value of AMPA synaptic weight *µ*_AMPA_. For a three-fold postsynaptic probability change (generated using simulated data), the shift in the mean value of AMPA synaptic weight is even more evident, suggesting a dose-dependent relationship between synaptic strength changes and AMPA receptor weight modulation (Figure 8Dii).

Correlation analysis between changes in postsynaptic spike probability and model parameters, sampled from the prior, revealed that while multiple parameters correlate with synaptic probability, it is most strongly correlated with the change in the mean value *µ*_*AMP A*_ of the AMPA receptor component in the model (Figure 8E). This predominant role of AMPA receptors aligns with established LTP mechanisms, where initial potentiation typically involves insertion of additional AMPA receptors into the postsynaptic membrane. Furthermore, as is evident in Figure 8F, the association between the two variables is nonlinear, suggesting that while initial increases in AMPA weight produce substantial enhancements in postsynaptic firing probability, the effect gradually saturates at higher AMPA values, consistent with physiological constraints on synaptic strengthening.

By analyzing the distributions of the MAP model parameter configurations of the neuronal pairs before and after 5 Hz stimulation (Figure 8 G), all parameters shown in Figure S12D), we observed significant changes across multiple synaptic components following stimulation. The most notable change occurs in the mean synaptic weight of the AMPA receptor (µ_*AMP A*_), whose distribution becomes right-skewed after stimulation. The increased right-skewness suggests that potentiation is confined to a subset of synapses, while the majority remain weak, rather than reflecting a uniform shift across the population. These findings align with biological studies of early-phase LTP [91], where AMPA receptor modulation precedes structural changes. The pre-LTP state involves a baseline complement of AMPA and NMDA receptors, while early LTP is characterized by rapid AMPA receptor insertion and an increase in the AMPA/NMDA ratio (Figure 8H). Interestingly, these changes were less pronounced when comparing pre-stimulation measurements with those taken 5-10 minutes after stimulation (Figure S12E), a period when postsynaptic spike probability is returning to baseline levels.

Some of the other parameters (*d*_*post*_, µ_*NMDA*_, *z*_*syn*_, 1/*λ*_*pre*_) also exhibited significant alterations following stimulation. These changes suggest more complex mechanisms related to the interplay of physiological and modeling factors. For example, the observed decrease in the NMDA receptor parameter following stimulation could be explained by several factors: First, in many synaptic plasticity models, the AMPA/NMDA ratio might be more critical than the absolute receptor values. Thus, the increase in AMPA receptor parameters may be accompanied by a relative decrease in NMDA parameters to reflect this balance rather than uniform increases. Second, synaptic changes during potentiation are not isolated events but involve coordinated modifications across multiple components, including changes in the composition and morphology of synapses, which may indirectly influence the summary statistics and the model”s interpretation of NMDA receptor contribution. Additionally, as seen in Figure 7E), NMDA shows prominent correlation with postsynaptic neuron firing rate, which changes significantly immediately upon stimulation (Figure S12A).

The ability to directly observe and quantify these mechanistic aspects of synaptic potentiation in isolated neuronal pairs demonstrates the analytical power of the presented platform for investigating fundamental synaptic plasticity mechanisms in a controlled environment.

## Discussion

*In vivo* neuronal circuits operate within highly complex environments with numerous confounding factors that often obscure fundamental synaptic mechanisms. The platform presented in this work reduces this complexity by providing a simplified environment where excitatory neurons form controlled connections in PDMS microstructures. This reductionist approach offers several advantages for investigating fundamental questions about synaptic function: it allows for direct observation of signal transmission between defined neuronal pairs without interference from surrounding networks, eliminates variables like heterogeneous cell types or inhibitory influences that typically complicated recordings, and enables precise manipulation of specific connections while maintaining long-term viability and recording capabilities.

We have achieved several key advances in studying human synaptic function with this platform. We successfully maintained hundreds of isolated parallel neuronal pairs for more than 100 days, outperforming other existing approaches[93, 20, 12, 77, 11, 94, 95, 19] both in terms of the recording throughput and the longevity of cultures, even without the presence of support cells. We showed that the NGN2 cultures are electrophysiologically active for several months *in vitro* despite being isolated. We further demonstrated that even the smallest network-level configuration — isolated neuronal pairs — exhibited functional synapses. The robust recovery of functional connectivity after washout of the synaptic blockers highlighted the system”s stability, making it particularly valuable for longitudinal studies of synaptic plasticity and pharmacological interventions. The system”s stability enabled us to collect large electrophysiological datasets, providing both detailed single-neuron information and collective population behavior. While plasticity mechanisms have remained largely a “black box” when investigated with extracellular electrophysiology alone in network-level cultures [96] and organoids [97] we addressed this limitation by developing a computational approach that enabled fitting large quantities of electrophysiological data without requiring extensive computational resources. The biophysical modeling framework demonstrated its power through the stimulation paradigm as an example application, where even modest experimental changes (*e.g*., 10% increases in synaptic probability) could be decomposed into specific underlying synaptic parameter modifications, revealing the sensitivity and utility of our integrated approach.

The main advantage of the platform is the relative simplicity of substrate preparation and use as opposed to other single-cell approaches that require extensive preparation steps [98, 93, 19]. The modularity in the design and dimensions as opposed to more standard microfluidic approaches[20, 99], can accommodate diverse experimental configurations ranging from studying basic connectivity to investigating complex plasticity phenomena, and from monocultures to co-cultures with defined pre- and postsynaptic populations. The possibility of integration with nanochannels [21, 18] allows for subcellular analysis. Unlike existing combined *in vitro*/*in silico* approaches that capture only overall culture behavior [100, 101, 33], the framework allows for the extraction of detailed statistics from specific neuronal pairs, the analysis of their properties through computational modeling, and the subsequent experimental investigation of the same pairs, creating a bidirectional feedback loop between modeling and experimentation at the single-neuron level. To our knowledge, this represents the first platform to integrate single-cell resolution with long-term culturing capabilities, an automated data analysis pipeline, biophysical modeling, and adaptable experimental design suitable for diverse cell types and experimental paradigms.

Nevertheless, several limitations of our current approach should be considered for future development. The stochastic nature of neuronal entry into microchannels meant we lacked precise control over the number of neurons within each structure, introducing variability in experimental conditions. Additionally, NGN2-derived excitatory neurons developed extensive axonal projections that could exceed the channel length limitations imposed by CMOS chip dimensions, potentially causing undesired backgrowth. While incorporating curved channel designs could accommodate longer axons, this modification would reduce the number of replicates per chip, thereby reducing statistical power.

The connectivity potentiation studies presented here, while demonstrating statistically significant changes in postsynaptic spike probability, represent only an initial step into the study of plasticity mechanisms. For more complex electrical stimulation paradigms [8, 102, 103], one would need to account for cell-electrode positions as well as conduction and synaptic delays between identified pre- and postsynaptic pairs in more detail. If this limitation is addressed, theoretical learning rules such as Bienenstock–Cooper–Munro (BCM) learning rule [104] or spike-time-dependent plasticity (STDP) [24, 25] along with its variations [103, 105] could be tested. Expanding the repertoire of stimulation frequencies and stimulation delays would thus be a promising next step. Additionally, the computational model could be enhanced with greater biophysical realism by accounting for receptor kinetics, calcium dynamics, and vesicle release mechanisms and/or with incorporating synaptic learning rules. This would open possibilities to generate and validate new hypotheses about how the potentiation occurs upon stimulation.

Despite the limitations mentioned above, the adaptability of our system supports numerous promising applications across neuroscience research and beyond. The platform serves as an optimal foundation for incrementally increasing system complexity in a controlled manner, where researchers could systematically introduce additional elements — such as inhibitory neurons, specific glial populations, or defined neuromodulators — and directly observe their effects on synaptic function and information transmission.

Additional applications include high-throughput drug screening with single-neuron resolution, investigation of synaptic transmission failures and their recovery mechanisms, and systematic characterization of disease-related synaptic dysfunction in patient-derived neurons. The platform”s compatibility with optical techniques also opens possibilities for combined electrophysiological and optical studies, enabling correlation between structural and functional synaptic changes. By enabling this step-by-step approach to increasing complexity, the system provides a powerful tool for testing specific hypotheses about fundamental mechanisms underlying synaptic function, plasticity, and network-level information processing, potentially helping to bridge the gap between simplified *in vitro* models and the intricate reality of *in vivo* neuronal circuits.

In conclusion, our platform represents a powerful tool for studying the human synaptic function, combining the precision of single-pair isolation with the statistical power of high-throughput analysis. The integration of PDMS microstructures, HD-MEAs, and biophysical modeling created a versatile experimental framework that bridges the gap between reductionist approaches and biological complexity. As neuroscience increasingly demands human-relevant models for understanding brain function and dysfunction, our platform offers a scalable solution for investigating the fundamental mechanisms of neuronal communication, plasticity, and network formation with high detail and throughput.

## Supporting information

AP propagation and transmission on CMOS

Cell migration on DIV1

## Acknowledgments

This work was supported by ETH Zurich and Swiss National Science Foundation (Project Nr. 182779). The authors thank Dr. Julian Hengsteler for video editing and compiling. The authors further thank other LBB members for fruitful discussions.

## Author Contributions

GA: Conceptualization, Methodology, Supervision, Software, Validation, Formal analysis, Investigation, Writing, Visualization. VV: Conceptualization, Methodology, Software, Validation, Formal analysis, Investigation, Writing, Visualization. JD: Conceptualization, Methodology, Supervision, Writing – review & editing. MLAS: Validation, Formal analysis, Software, Investigation, Writing – review & editing. TS: Validation, Formal analysis, Investigation, Writing – review & editing. AS: Validation, Formal analysis, Investigation, Writing – review & editing. FCT: Validation, Formal analysis, Software, Investigation, Writing – review & editing. JK: Validation, Formal analysis, Software, Investigation, Writing – review & editing. TR: Investigation, Writing – review & editing. JV: Conceptualization, Resources, Project administration, Funding acquisition, Writing – review & editing. KV: Conceptualization, Methodology, Supervision, Software, Validation, Formal analysis, Investigation, Writing, Visualization.

## Generative AI statement

The author(s) declare that Generative AI was used in the creation of this manuscript. Generative AI was used for language edits.

## Data Availability

The code used in this publication will be available on GitLab. Data will be available through ETH Research Collection. Supplemwntary videos are available on this link.

## Footnotes

1 *ALaboratory of Biosensors and Bioelectronics (LBB), Institute for Biomedical Engineering, ETH Zurich, Zurich, Switzerland* ^*∗*^ *Corresponding author*

