## Supplementary Figures and Tables for "An Integrated *In Vitro* Platform and Biophysical Modeling Approach for Studying Synaptic Transmission in Isolated Neuronal Pairs"

### Supplementary Material

#### Supplementary Figures

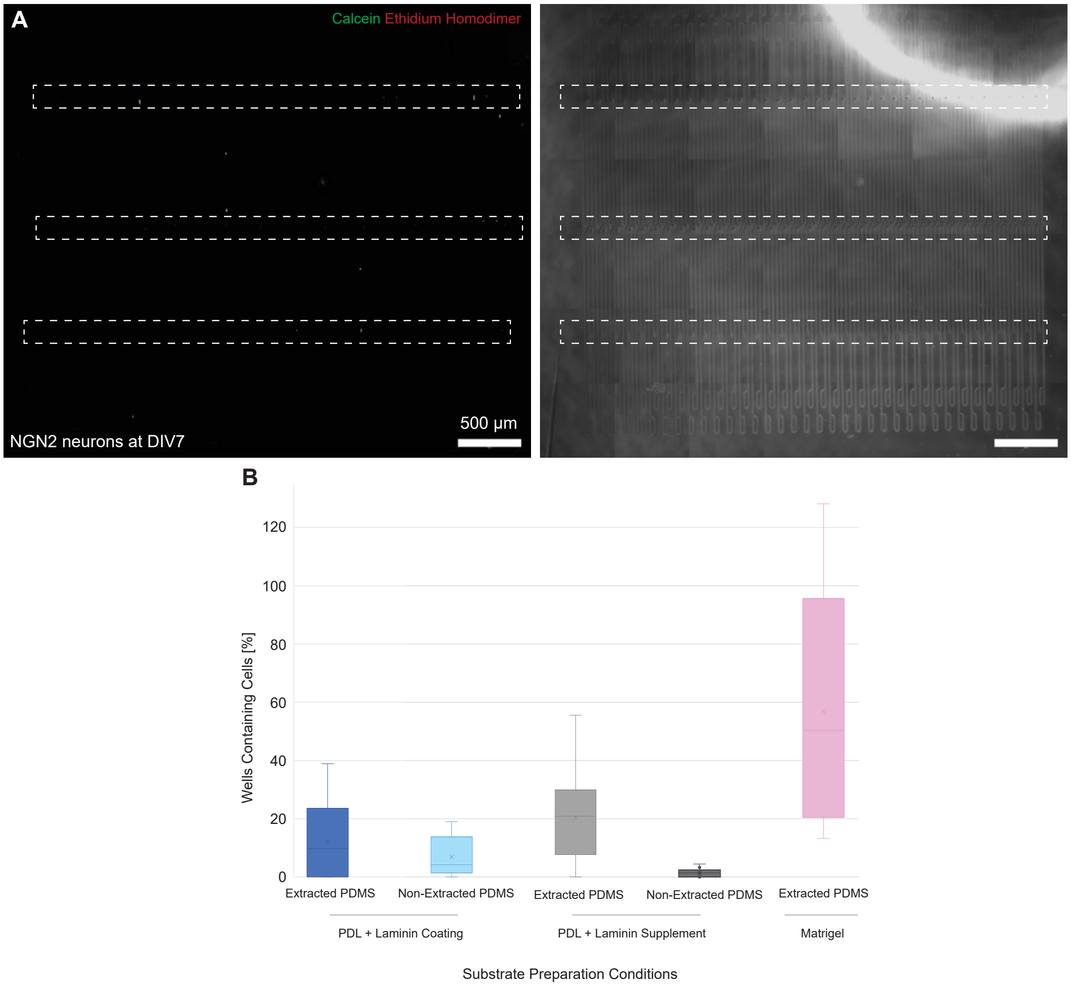


**Fig. S1**: **A**) Microscopy image showing low number of isolated cells on DIV7 inside a microstructure placed on the substrate coated with PDL and laminin. The culture was stained with live/dead staining kit consisting of calcein and ethidium homodimer, which is shown on the left. The phase contrast image showing microchannel outline is shown on the right. Well area is highlighted with white dashed squares. **B**) Comparisons of cell abundance inside the microstructures for different substrate preparations. Following cell-adhesive coating conditions are compared: PDL coating and laminin added as a coating before microstructure mounting, PDL coating and laminin added as a medium supplement after microstructure mounting and matrigel as a coating added after microstructure mounting. Extracted and non-extracted PDMS microstructures are compared for the first two conditions.

#### Supplementary Tables

Table S1: Kruskal-Wallis independent samples test results comparing synaptic probability across different experimental timepoints: before adding the synaptic blocker, immediately after adding the blocker and one-week-after upon washout. Statistical significance is denoted as: ns ($p > 0.05$), * ($0.01 < p \leq 0.05$), ** ($0.001 < p \leq 0.01$), *** ($0.0001 < p \leq 0.001$), and **** ($p \leq 0.0001$).

| **Metric** | **Before vs. Blocked** | **Blocked vs. Washed** | **Before vs. Washed** |
|---|---|---|---|
| Postsynaptic Spike Probability | $P=8.388 \cdot 10^{-9}$<br>Stat=33.18<br>**** | $P=3.197 \cdot 10^{-10}$<br>Stat=39.55<br>**** | $P=9.517 \cdot 10^{-1}$<br>$\text{Stat}=3.672 \cdot 10^{-3}$<br>ns |

Table S2: Kruskal-Wallis independent samples test results comparing connectivity metrics before and after stimulation as shown in Fig. 8. B represents baseline recording prior to stimulation, A represents the first 5 minutes post-stimulation, and $A_2$ represents the second 5 minutes post-stimulation. Statistical significance is denoted as: ns ($p > 0.05$), * ($0.01 < p \leq 0.05$), ** ($0.001 < p \leq 0.01$), *** ($0.0001 < p \leq 0.001$), and **** ($p \leq 0.0001$).

| **Metric** | **B vs. A** | **A vs. $A_2$** | **B vs. $A_2$** |
|---|---|---|---|
| Postsynaptic Spike Probability | P=0.01<br>Stat=6.224<br>* | P=0.3<br>Stat=1.314<br>ns | P=0.1<br>Stat=2.540<br>ns |
| mTE | P=0.002<br>Stat=9.368<br>** | P=0.06<br>Stat=3.519<br>ns | P=0.2<br>Stat=2.059<br>ns |
| Lag | P=0.6<br>Stat=0.246<br>ns | P=0.2<br>Stat=1.615<br>ns | P=0.08<br>Stat=3.049<br>ns |

Table S3: Statistical analysis of HH model parameters comparing experimental data before and immediately after stimulation. Three different tests were applied: t-test (tests if the means of the distributions are different), KS-test (tests if the entire distributions are different), and Mann-Whitney U test (non-parametric test for distribution differences).

| **Parameter** | **t-test** | **Sig.** | **KS-test** | **Sig.** | **MW-test** | **Sig.** |
|---|---|---|---|---|---|---|
| $d_{pre}$ | P= 0.9 | False | P=0.5 | False | P=0.8 | False |
| $d_{post}$ | P=0.02 | True | P=0.03 | True | P=0.04 | True |
| $mean_{AMPA}$ | P=0.003 | True | P=0.008 | True | P=0.001 | True |
| $std_{AMPA}$ | P=0.1 | False | P=0.03 | True | P=0.2 | False |
| $mean_{NMDA}$ | P=0.006 | True | P=0.02 | True | P=0.01 | True |
| $std_{NMDA}$ | P=0.8 | False | P=0.2 | False | P=0.3 | False |
| $1/\lambda_{pre}$ | P=0.03 | True | P=0.0006 | True | P=0.09 | False |
| $1/\lambda_{post}$ | P=0.1 | False | P=0.01 | True | P=0.09 | False |
| Syn. Loc. | P=0.03 | True | P=0.01 | True | P=0.03 | True |

Table S4: Statistical analysis of HH model parameters comparing experimental data before and five minutes after stimulation. Three different tests were applied: t-test (tests if the means of the distributions are different), KS-test (tests if the entire distributions are different), and Mann-Whitney test (non-parametric test for distribution differences).

| **Parameter** | **t-test** | **Sig.** | **KS-test** | **Sig.** | **MW-test** | **Sig.** |
|---|---|---|---|---|---|---|
| $d_{pre}$ | P=0.6 | False | P=0.2 | False | P=0.8 | False |
| $d_{post}$ | P=0.08 | False | P=0.08 | False | P=0.2 | False |
| $mean_{AMPA}$ | P=0.07 | False | P=0.01 | True | P=0.007 | True |
| $std_{AMPA}$ | P=0.08 | False | P=0.5 | False | P=0.4 | False |
| $mean_{NMDA}$ | P=0.001 | True | P=0.02 | True | P=0.007 | True |
| $std_{NMDA}$ | P=0.6 | False | P=0.8 | False | P=0.9 | False |
| $1/\lambda_{pre}$ | P=0.3 | False | P=0.1 | False | P=0.4 | False |
| $1/\lambda_{post}$ | P=0.5 | False | P=0.5 | False | P=0.8 | False |
| Syn Loc. | P=0.0007 | True | P=0.0003 | True | P=0.0004 | True |

Table S5: Mann-Whitney-Wilcoxon test results comparing non-normalized absolute deviation of parameters pulled from Maximum A Posterior (MAP) and Low Probability (LP) distributions. Statistical significance is denoted as: ns (p ¿ 0.05), * (0.01 < p ≤ 0.05), ** (0.001 < p ≤ 0.01), *** (0.0001 < p ≤ 0.001), and **** (p ≤ 0.0001).

| **Metric** | **Comparison** | **P-value** | **U-statistic** |
|---|---|---|---|
| Firing Rate | $Target_{MAP}$ vs. $Target_{LP}$ | $5.067 \cdot 10^{-150}$ **** | $3.311 \cdot 10^5$ |
| Firing Rate | $Source_{MAP}$ vs. $Source_{LP}$ | $1.492 \cdot 10^{-58}$ **** | $5.193 \cdot 10^5$ |
| Conduction Speed | $Source_{MAP}$ vs. $Source_{LP}$ | $8.346 \cdot 10^{-94}$ **** | $4.359 \cdot 10^5$ |
| Conduction Speed | $Target_{MAP}$ vs. $Target_{LP}$ | $3.393 \cdot 10^{-159}$ **** | $3.160 \cdot 10^5$ |
| Synaptic Probability | MAP vs. LP | $5.189 \cdot 10^{-146}$ **** | $3.378 \cdot 10^5$ |
| Lag | MAP vs. LP | $9.610 \cdot 10^{-115}$ **** | $4.108 \cdot 10^5$ |

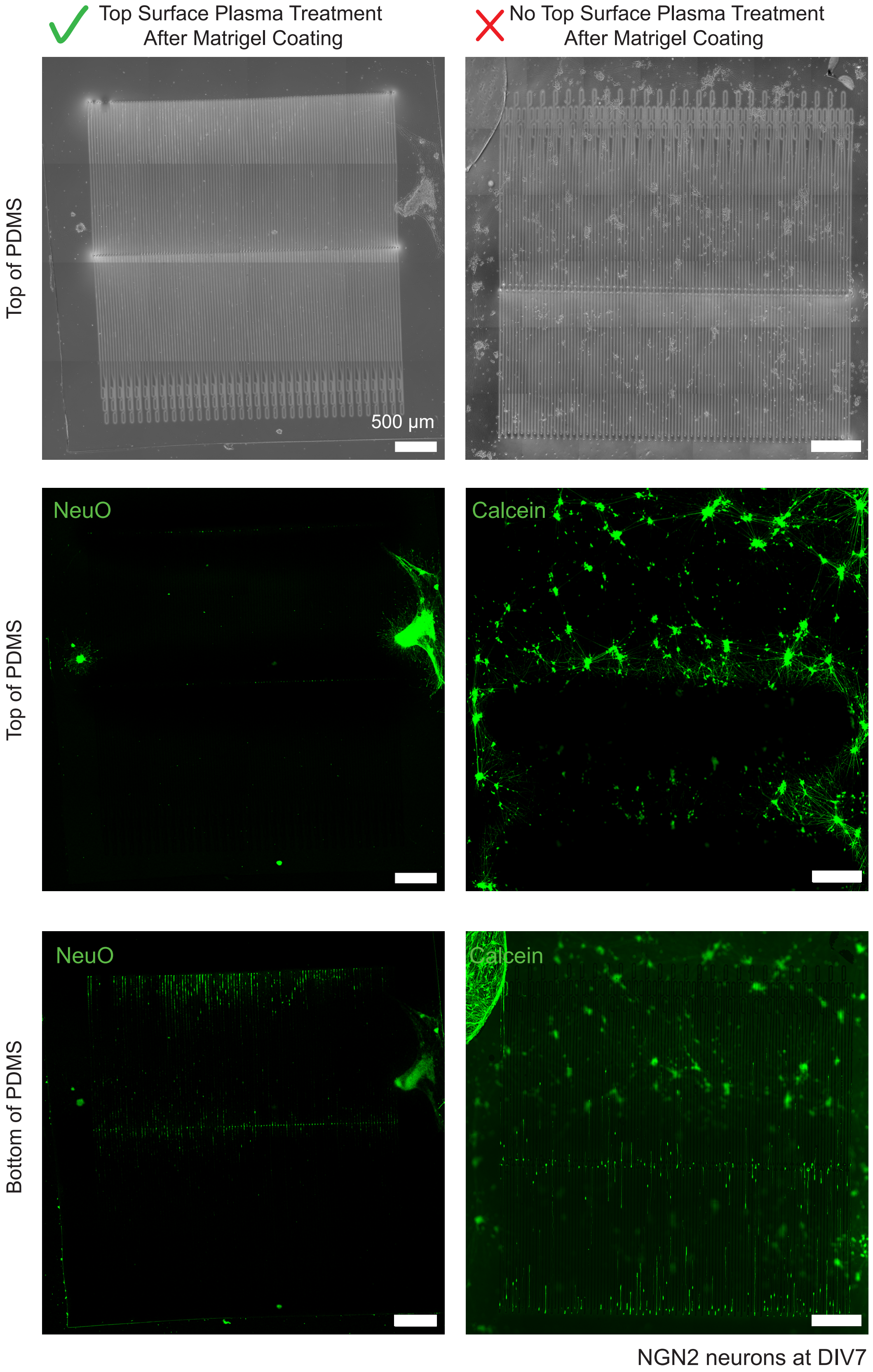


**Fig. S2**: Phase contrast (top raw) and fluorescent microscopy images showing the comparison of the PDMS top surface when treating the surface with plasma after matrigel coating (top left, middle left) and when no treatment was applied upon coating (top right, middle right). The abundance of cells inside the microchannels is comparable in both cases (bottom left and right), indicating that matrigel coating inside the microchannels is preserved upon plasma treatment. All images were obtained on DIV 7.

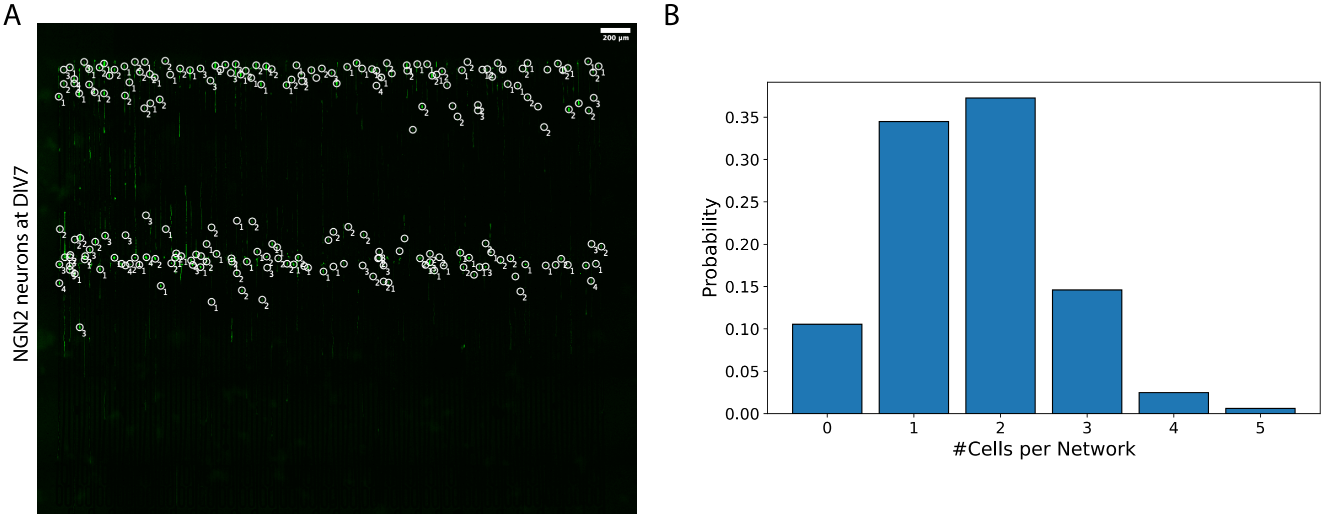


**Fig. S3**: Quantification of cell counts per network. A) Fluorescence microscopy image of a microstructure containing multiple cells. Individual somata were manually identified and annotated; somata located within the same network were grouped to obtain the cell count per network. B) Distribution of the number of cells per network across all analyzed networks (n=322).

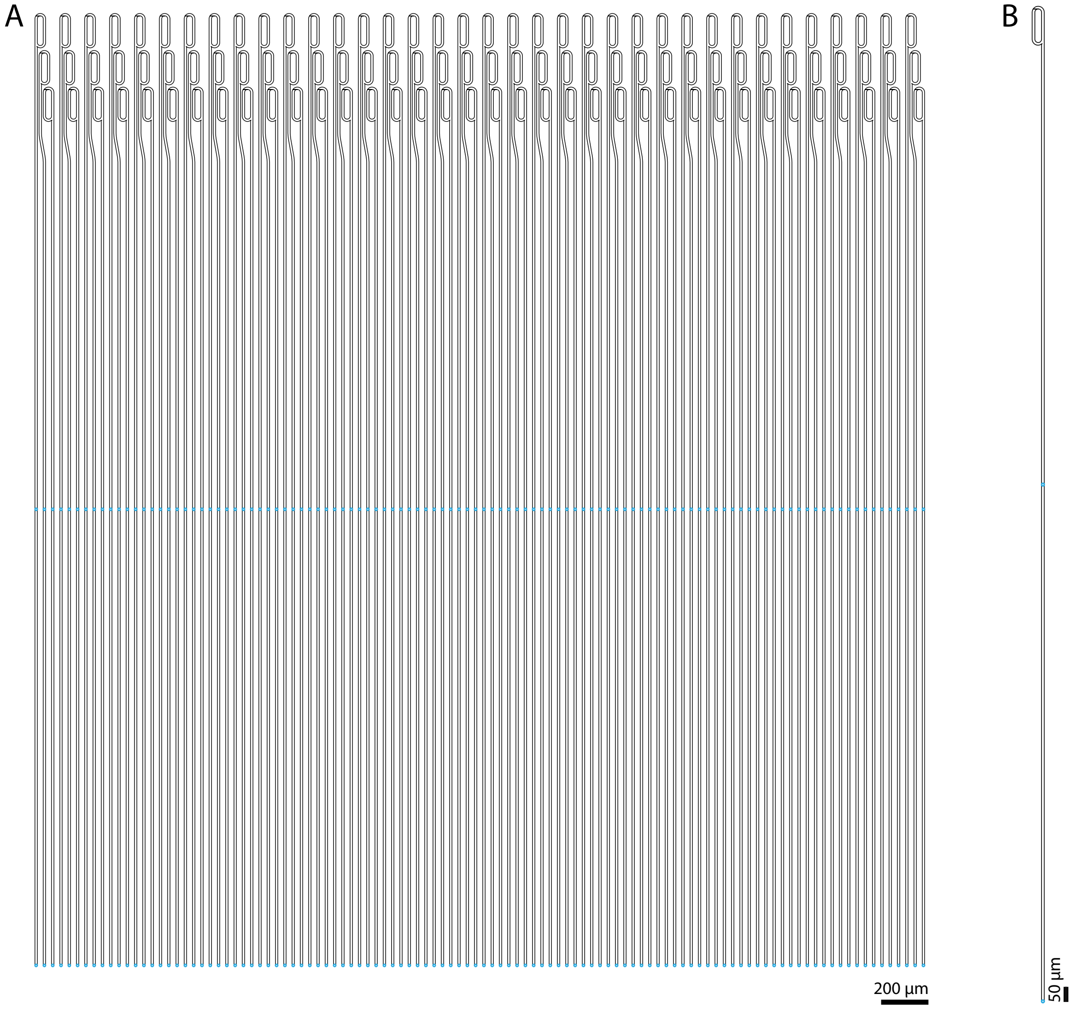


**Fig. S4**: Overview of a straight microstructure design. A) One microstructure that fits on a HD-CMOS MEA contains 108 isolated microchannels with two rows of 10 µm openings (light blue dots). B) Single microchannel example.

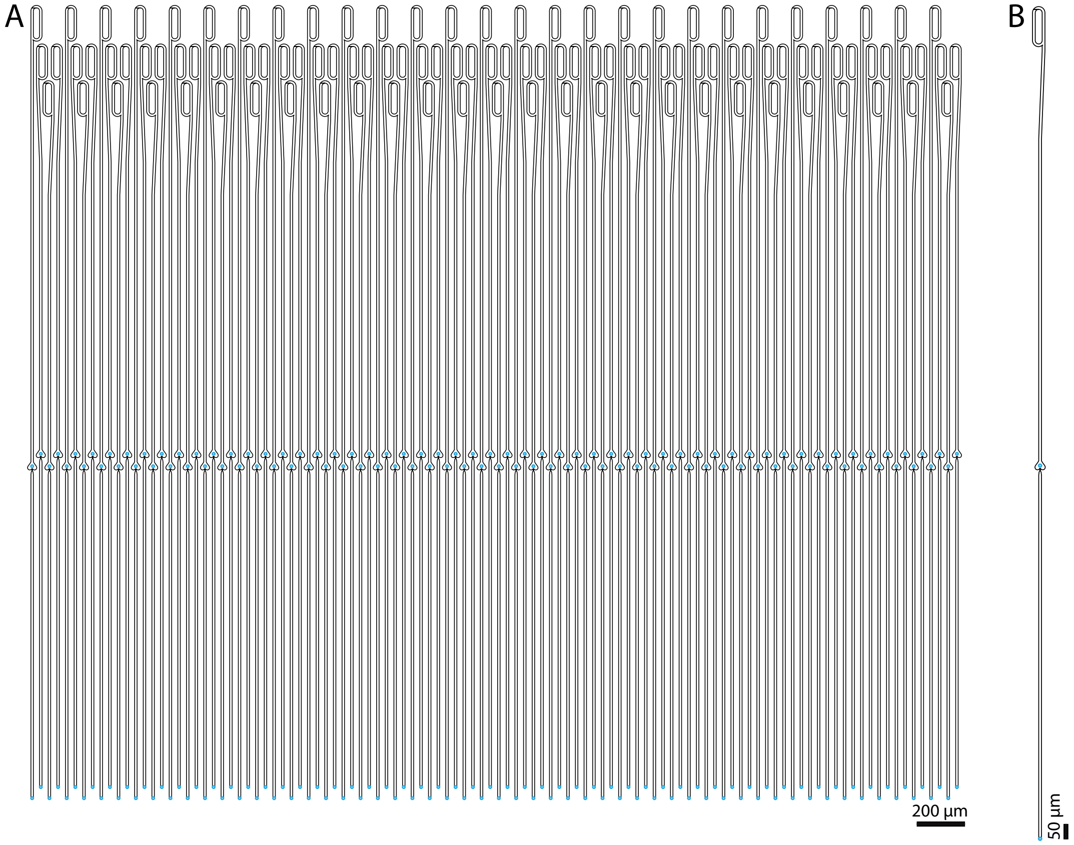


**Fig. S5**: Overview of a heart microstructure design. The heart shape promotes preferential axon growth towards its apex. A) One microstructure that fits on a HD-CMOS MEA contains 108 isolated microchannels with two rows of 10 μm openings (light blue dots). B) Single microchannel example.

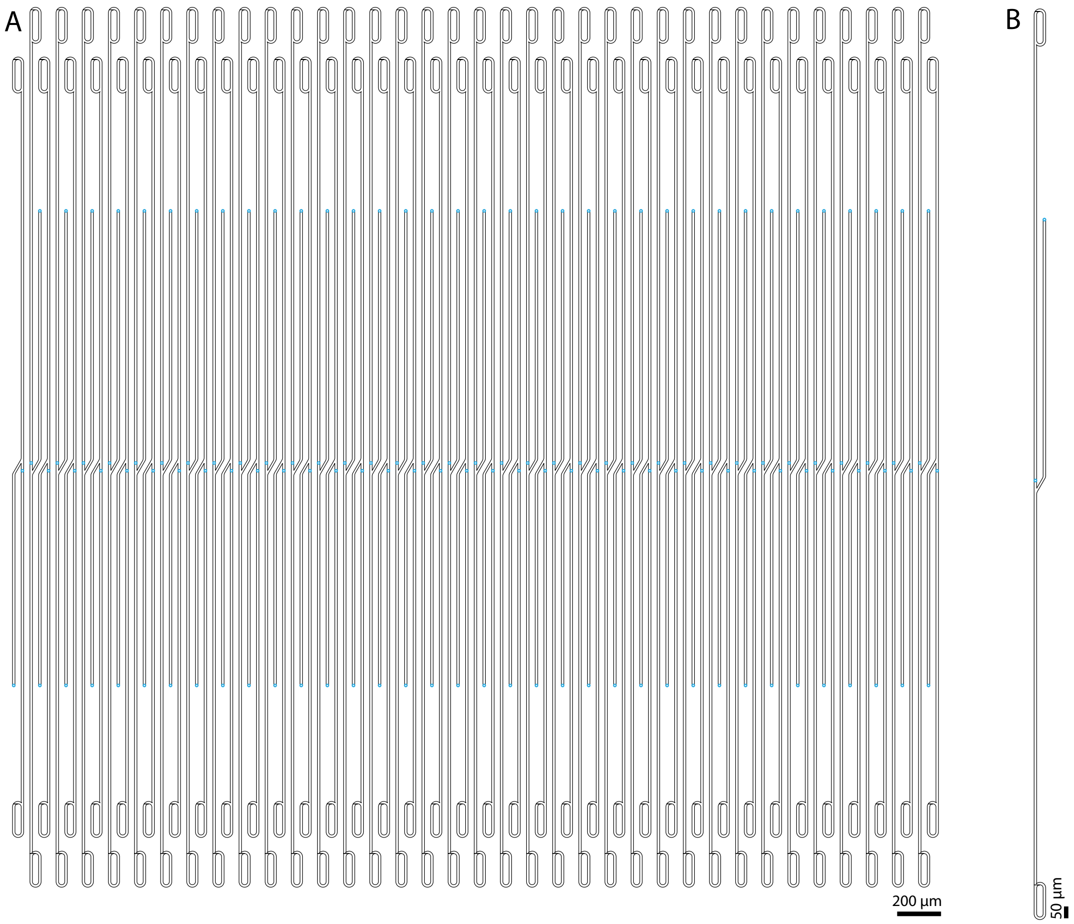


**Fig. S6**: Overview of an en-passant microstructure design. This geometry assures that the axon of the presynaptic neuron grows close enough to connect to the dendrites of the postsynaptic cell. A) One microstructure that fits on a HD-CMOS MEA contains 72 isolated microchannels with three rows of 10 µm openings (light blue dots). B) Single microchannel example.

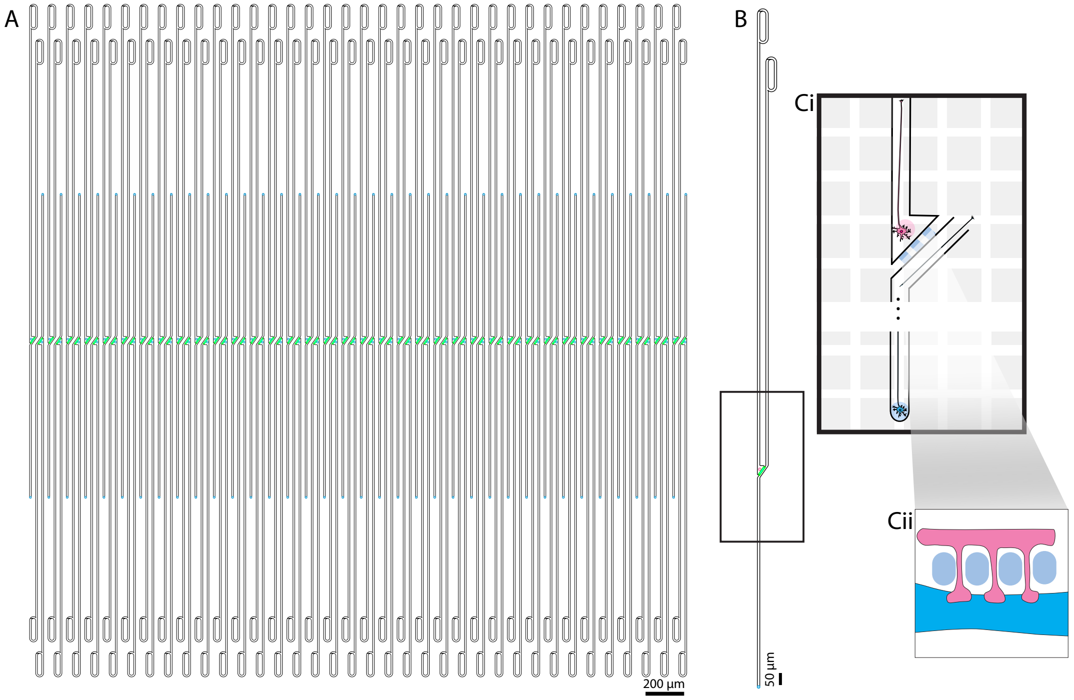


**Fig. S7**: Overview of an 1-in-1-out nanochannel microstructure design. Here, nanochannels prevent axons to grow through but enable connection through synaptic spines [21]. A) One microstructure that fits on a HD-CMOS MEA contains 72 isolated microchannels with three rows of 10 µm openings (light blue dots). B) Single microchannel example.

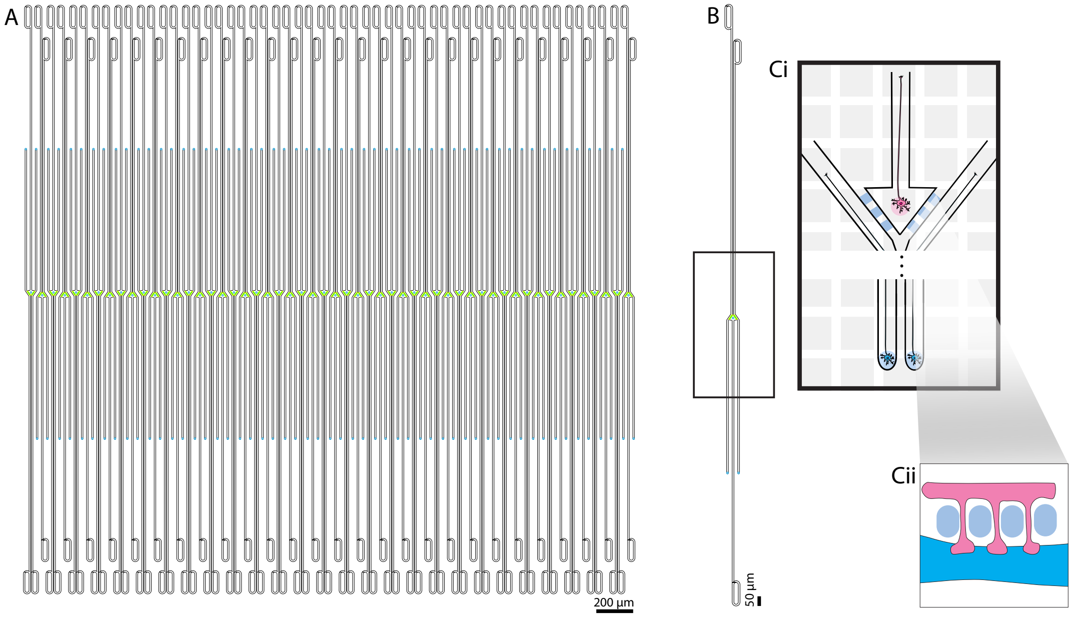


**Fig. S8**: Overview of a 2-in-1-out nanochannel microstructure design that enables connecting two axons to the dendrites of a postsynaptic neuron. A) One microstructure that fits on a HD-CMOS MEA contains 44 isolated microchannels with three rows of 10 µm openings (green dots indicating the position of the postsynaptic, light blue dots the presynaptic neurons). B) Single microchannel example.

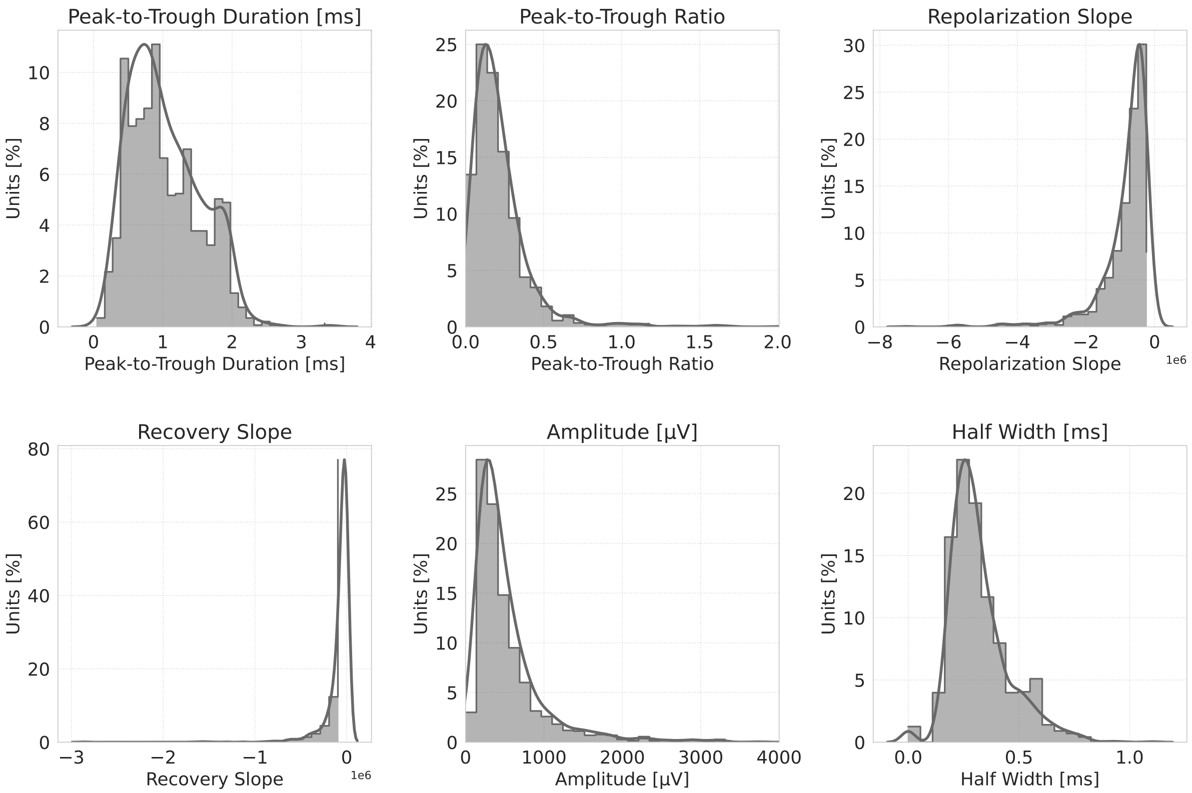


**Fig. S9**: Waveform metrics extracted by custom-designed data analysis pipeline after the spike sorting performed using the Spikeinterface framework.

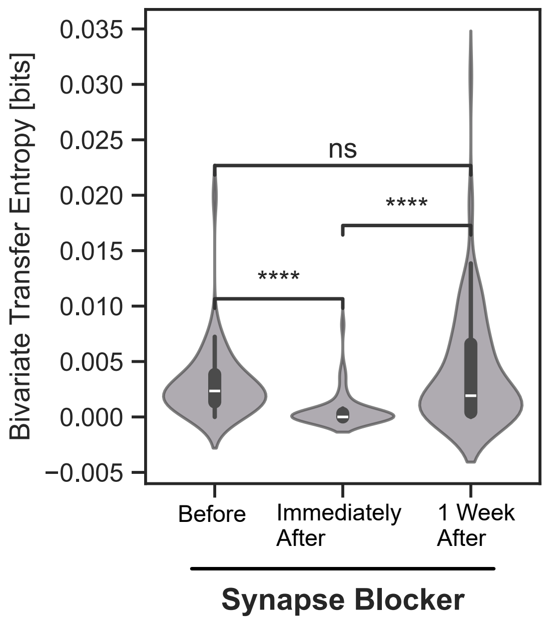


**Fig. S10**: Bivariate transfer entropy calculated for recordings before, immediately after and one week after adding synaptic receptor antagonists. Statistical test details are available in Table S1.

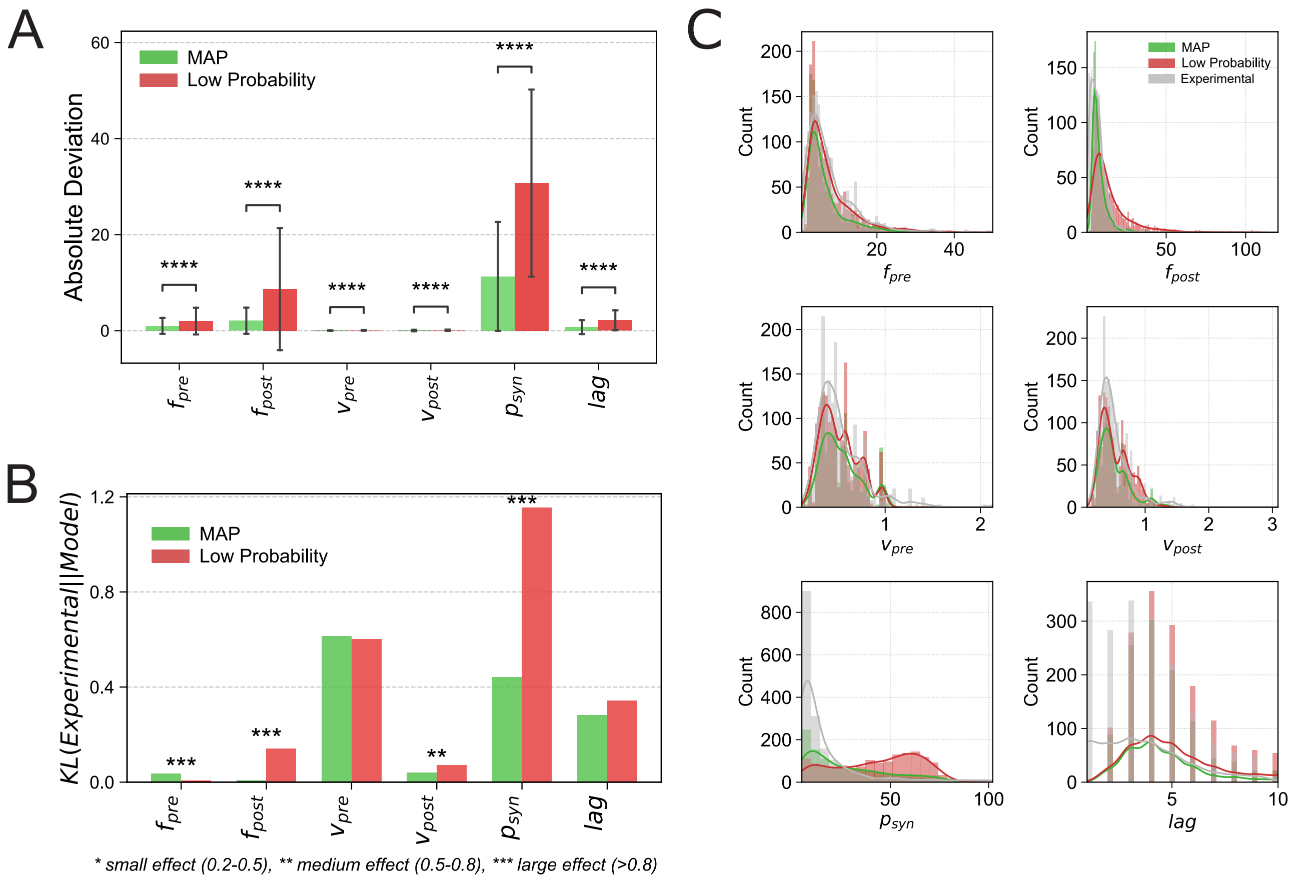


**Fig. S11**: **A**) Absolute deviation between parameters sampled from high-probability (maximum a posteriori) and low-probability regions of the distribution. **B**) Kullback-Leibler divergence comparison between maximum a posteriori and low-probability parameter sets. **C**) Visual comparison of maximum a posteriori, low-probability, and experimentally observed parameters.

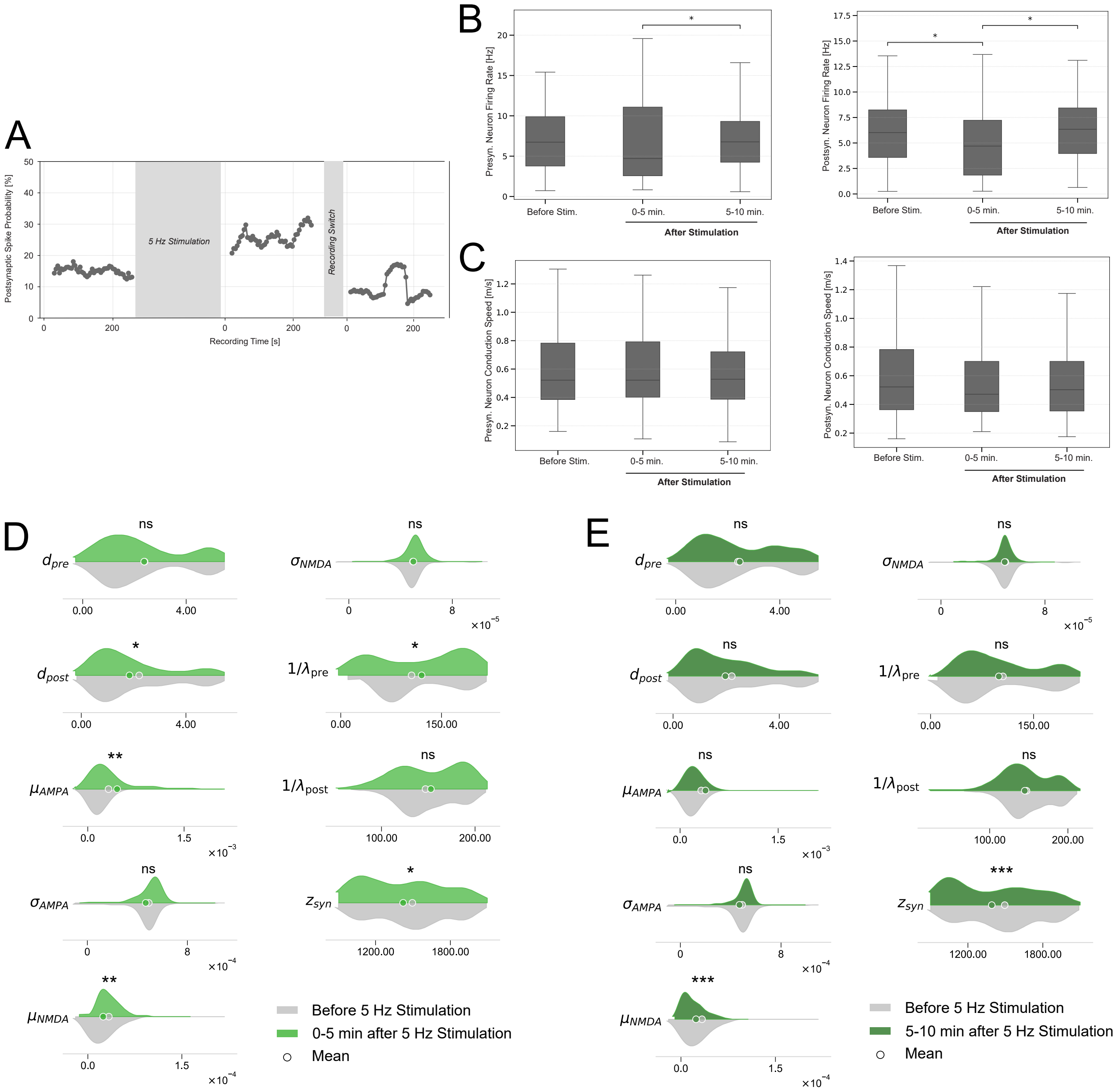


**Fig. S12**: **A** Mean postsynaptic spike probability over time with corresponding confidence intervals, concatenated across two post-stimulation recording sessions. **B**) Firing rate per neuron extracted from recordings before, immediately after and five minutes after the stimulation protocol was applied. Left plot captures presynaptic neurons (sources), and right plot captures postsynaptic neurons (targets). **C**) Conduction speed per neuron extracted from recordings before, immediately after and five minutes after the stimulation protocol was applied. Left plot captures presynaptic neurons (sources), and right plot captures postsynaptic neurons (targets). **D**) Distribution comparison for parameters inferred by the model from recordings before and immediately after stimulation. Significantly different distributions are highlighted in red. **E**) Distribution comparison for parameters inferred by the model from recordings before and five minutes after stimulation. Significantly different distributions are highlighted in red.

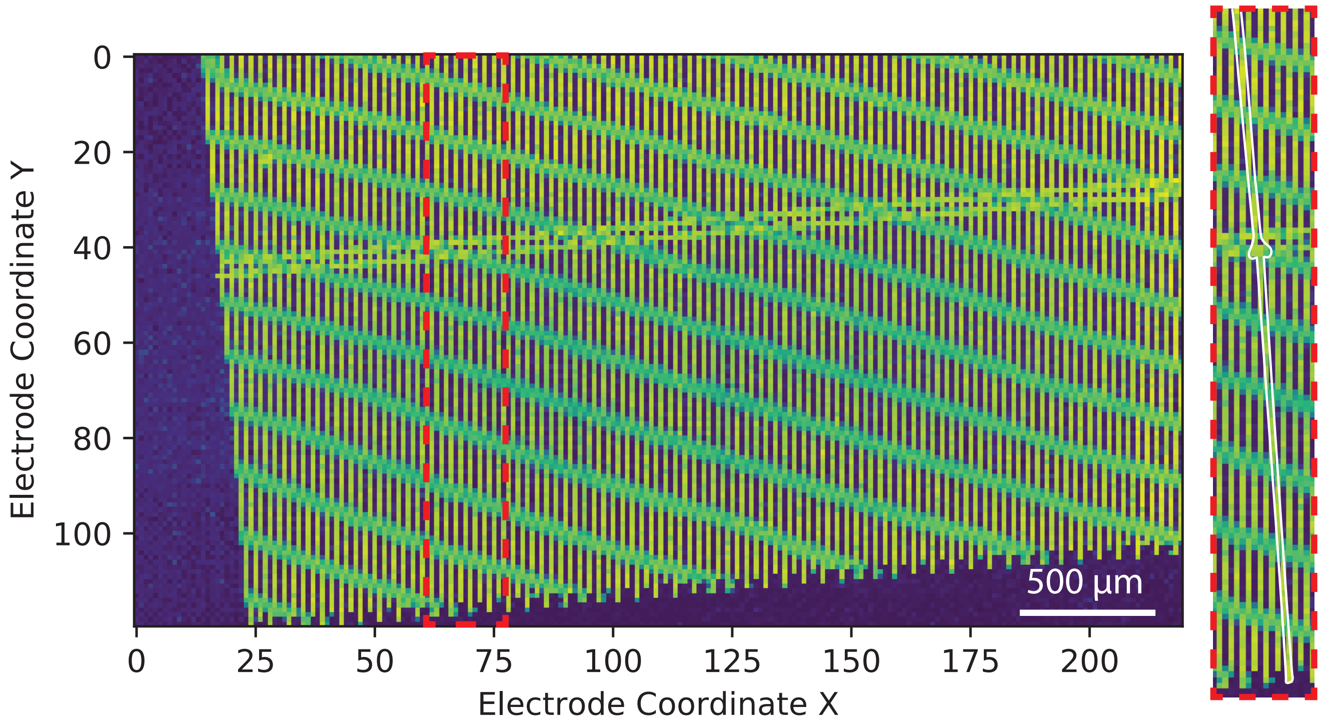


**Fig. S13**: An example of a voltage map obtained after microstructure adhesion to the chip. The difference in impedance between electrodes that are covered in PDMS and that are exposed to the liquid enables us to identify the independent networks.

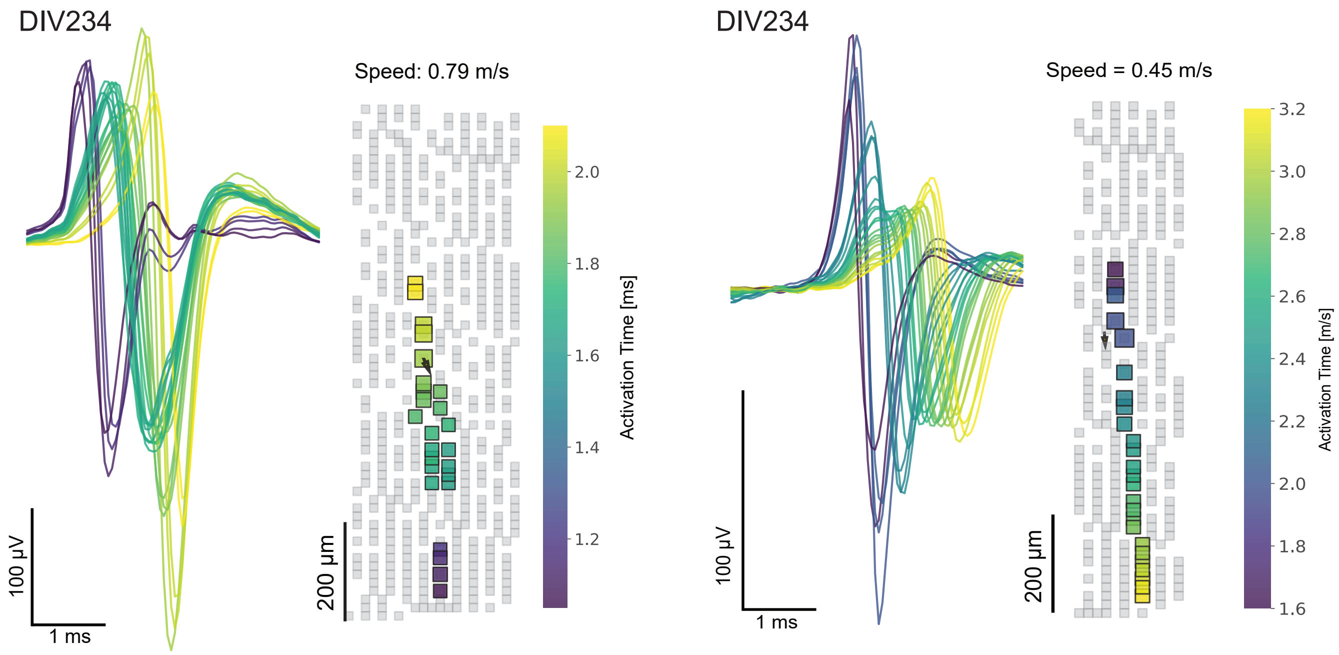


**Fig. S14**: Waveforms extracted from DIV 234 recordings from NGN2 neurons isolated in microchannels.

#### Prediction of Firing Probability with Machine Learning Models

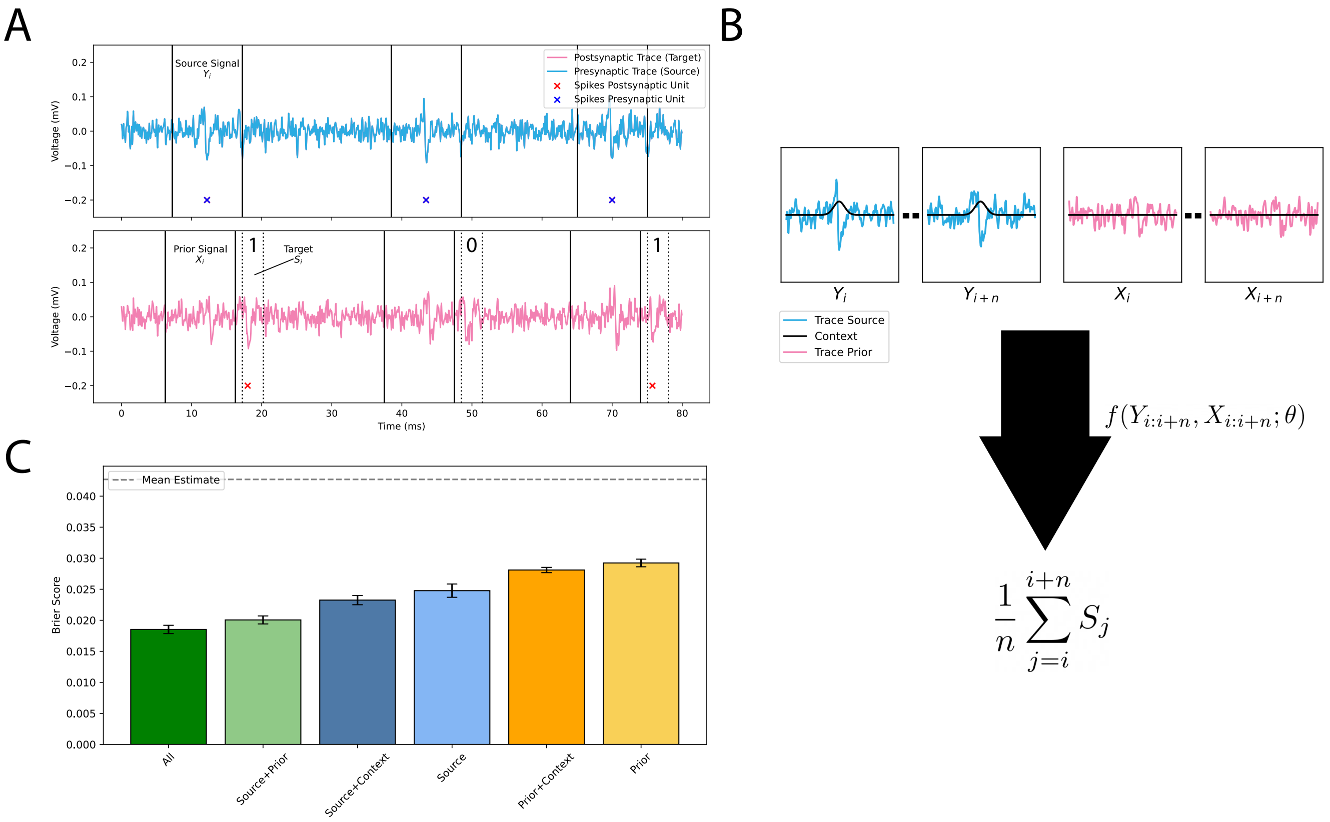


**Fig. S15**: **A**) Illustration of sample extraction from raw extracellular signals. The target window (3 ms) is centered around the most significant lag derived from the transfer entropy between pre- and postsynaptic neuron pairs. Source and prior time windows are aligned relative to the target. The binary target is set to one if a postsynaptic spike occurs within the window, zero otherwise. **B**) Machine learning setup: a transformer model receives $n$ samples and predicts the expected firing probability. An optional context signal (spike train convolved with a Gaussian) can be added. **C**) Brier scores for various model configurations. A mean baseline, which predicts the average firing probability from the training set, is included as a reference.

To predict whether a postsynaptic spike occurs within a defined time window, we trained a transformer model $f(\cdot;\theta)$ on cutouts of extracellular voltage traces. The aim of this analysis is to evaluate whether the neuron pairs identified through the transfer entropy approach carry sufficient information in their prior and source (presynaptic neuron) signals to enable reliable prediction of postsynaptic spiking probability. This serves two purposes: first, to validate that the variables used in the transfer entropy analysis indeed reflect a directional information exchange—without which a predictive mapping would be impossible; and second, to gain insight into which signal components are more informative for postsynaptic spiking, thereby shedding light on the temporal structure of the underlying interaction. Pre- and postsynaptic neuron pairs were selected such that each neuron's extremum electrode appeared only once in the dataset. The timing of events for sample extraction was based on spike times of the presynaptic neuron, adjusted by the most significant lag obtained from the transfer entropy analysis:

$$e_i^k = s_{\text{pre},i}^k + \text{lag}_k \tag{7}$$

where $s_{\text{pre},i}^k$ is the $i$-th spike time of the presynaptic neuron in pair $k$, and $\text{lag}_k$ is the transfer entropy lag for that pair.

From each event time $e_i^k$, fixed-length signal snippets are extracted:

$$\begin{aligned} Y_i[t] &= R_{pre}^k[e_i^k + t - \Delta_{\text{source}}], \quad t \in [0, w_{\text{source}}] \\ X_i[t] &= R_{post}^k[e_i^k + t - \Delta_{\text{prior}}], \quad t \in [0, w_{\text{prior}}] \end{aligned} \tag{8}$$

Here, $R^k$ denotes the extracellular recording from the corresponding extremum electrode, and $\Delta$ and $w$ denote the shift and window size for each signal. Context signals for $Y_i$ and $X_i$ are generated by selecting spike times within the same windows and convolving them with a Gaussian kernel (standard deviation 0.5 ms). The binary target $S_i$ is set to one if a spike from the postsynaptic neuron occurs within a bin of width $b$ centered at $e_i^k$, and zero otherwise. To form the input to the model, $n$ samples are concatenated, and the model is trained to predict the mean of the corresponding $n$ target values. Data was split into training (70%), validation (15%), and testing (15%) sets. Prior to the split, all input-target pairs were randomly permuted. Precise parameter values are given in Table S6.

The methodology and results are summarized in Figure S15. The model consistently outperforms a baseline predictor that uses the mean firing probability from the training data. While the model's performance is not perfect-as extracellular recordings, though informative, do not capture synaptic currents or the inherent noise in neuronal firing-the results clearly demonstrate that both source and prior signals contain predictive information about the postsynaptic spike. Among these, the source signal contributes more significantly, suggesting a stronger influence on the postsynaptic neuron's firing probability.

Table S6: Values used in the dataset creation for the machine learning approach.

| **Symbol** | **Value** | **Description** |
|---|---|---|
| $\Delta_{\text{source}}$ | 11 ms | Time shift applied to source window |
| $\Delta_{\text{prior}}$ | 12 ms | Time shift applied to prior window |
| $w_{\text{source}}$ | 10 ms | Duration of source window |
| $w_{\text{prior}}$ | 10 ms | Duration of prior window |
| $b$ | 3 ms | Bin width for target |
| $n$ | 10 | Number of samples concatenated as input |
